# APR-246 and DNA polymerase theta inhibitor Novobiocin synergize to eliminate *TP53* mutant AML cells

**DOI:** 10.64898/2026.10.30.763043

**Authors:** Amos Tuval, Terra Lasho, Anna Spada, Mrinal M. Patnaik, Klas G. Wiman

## Abstract

Patients with *TP53*-mutant myelodysplasic syndromes (MDS) and acute myeloid leukemia (AML) are usually elderly and frail with highly resistant malignant clones and a median survival of less than one year. The molecule APR-246 targets mutant p53 but also perturbs redox homeostasis. APR-246 showed favorable safety in a phase 3 trial in *TP53* mutant MDS/AML but did not meet efficacy endpoints in combination with azacitidine (AZA), highlighting the need for more effective combination strategies. Conversion of APR-246 to the active compound MQ releases formaldehyde which generates DNA-protein crosslinks. AML cells depend on DNA polymerase theta (PolƟ) to repair such lesions and PolƟ protects cells from ferroptosis. Using a panel of isogenic MOLM13 AML cell lines each carrying a common *TP53* variant, we show that the PolƟ inhibitor Novobiocin (NVB) synergizes with APR-246 to induce AML cell death. We demonstrate strong synergy with this combination in all *TP53* mutant cell lines and to a lesser extent in the *TP53* wildtype and null cells, indicating both p53-dependent and p53-independent mechanisms behind the synergy. APR-246 provides selectivity for *TP53*-mutated cells while NVB enhances p53-indendent apoptosis and p53-dependent ferroptosis. Levels of ferroptosis induction correlate with the observed synergy in *TP53*-mutant and WT cells. Combination of APR-246 and NVB induces potent *in vitro* growth inhibition of primary *TP53*-mutated AML/CMML patient samples, with significant activity achieved at low micromolar or nanomolar concentrations of APR-246. Our findings hold promise for combination of APR-246 and NVB as a new therapeutic strategy tailored for *TP53*-mutated AML patients with high efficacy and low toxicity.

## Introduction

Patients diagnosed with *TP53*-mutated MDS or AML have a median survival of <1 year (Shah et al. 2025a) and pose two main challenges: 1) These individuals are older in age, have co-morbidities and are often unable to tolerate intensive chemotherapy, including stem cell transplants (Shah et al. 2025b). 2) In most cases, malignant clones are characterized by biallelic *TP53* alterations (“multi-hit”) (Bernard et al. 2020), eliminating wildtype *TP53* function/activity, resulting in resistance to chemotherapy and currently available targeted agents for MDS/AML (Döhner et al. 2024). The combination of azacitidine or decitabine and venetoclax, currently considered the standard-of-care for patients that have adverse risk AML or who are ineligible for intensive chemotherapy, demonstrated a median overall survival of 5 months among *TP53*-mutated AML patients (Döhner et al. 2024; Kim et al. 2021). New combination treatments are being constantly sought for this patient subgroup, for example venetoclax with cladribine and low-dose cytarabine alternating with azacitidine that was specifically designed to reduce toxicity and limit outgrowth of resistant clones (Kadia et al. 2026, Amitai et al. 2026); however, the implementation and clinical outcomes of this combination were limited in *TP53*-mutated AML patients due to the above-mentioned challenges.

The compound APR-246 (eprenetapopt) is spontaneously converted to its active metabolite methylene quinuclidinone (MQ) which binds to several cysteines in p53, enhances p53 core domain thermostability, and induces tumor cell death (Lambert et al. 2009; Zhang et al. 2018; Degtjarik et al. 2021). MQ also has p53-independent effects, including glutathione depletion and induction of oxidative stress (Peng et al., 2013; Tessoulain et al. 2014; Liu et al. 2017; Ceder et al. 2021; Fujihara et al. 2022). APR-246 in combination with azacitidine (AZA) demonstrated encouraging clinical activity and a favorable safety profile in phase II trials in patients with *TP53*-mutated MDS/AML. A majority of the patients received more than 3 treatment cycles, with incidence rates and grades of myelosuppression that were not greater than those typically observed in patients receiving azacitidine monotherapy and neurological toxicities that were fully reversible following drug discontinuation, with no recurrence after appropriate dose modifications (Cluzeau et al. 2021; Sallman et al. 2021). However, although a phase III trial with the combination of APR-246 and AZA in *TP53*-mutated MDS/AML demonstrated a 50% increase in complete remission (CR) rates compared with azacitidine monotherapy (34.6% vs. 22.4%) (clinicaltrials.gov; NCT03745716), it did not meet efficacy endpoints (Aprea Therapeutics; www.aprea.com), indicating that novel strategies are needed to enhance its therapeutic activity.

One possible way of enhancing the efficacy of APR-246 is synergistic combination treatment with other agents. Previous studies have demonstrated synergy between APR-246 and DNA-damaging chemotherapeutic agents (Mohell et al. 2015), sulfasalazine that inhibits the cystine/glutamate antiporter xCT (Liu et al., 2017), PI3K/mTOR inhibitors that perturb anti-oxidant cellular protective mechanisms (Ali et al. 2016), and inhibitors of the XPO1 nuclear export protein (Kruer et al. 2025). Conversion of APR-246 to MQ releases formaldehyde (Lambert et al. 2009) that generates DNA-protein crosslinks (DPCs) (Vekariya et al. 2023). These DNA lesions can trigger p53-dependent hematopoietic stem cell attrition (Wang et al. 2023), but this has not been specifically exploited therapeutically thus far.

Novobiocin (NVB) is an aminocoumarin antibiotic that inhibits bacterial DNA synthesis by blocking the ATPase activity of DNA gyrase. It has been used for treatment of gram-positive bacterial infections at doses of 1-2 gr/day. Main toxicities at this dose range were allergic reactions and rash. Combination of NVB with chemotherapy in breast cancer patients in doses up to 5 gr per day did not cause increased hematological toxicity (Kennedy et al. 1995). NVB is currently being tested in patients with solid tumors and homology directed DNA repair deficiencies (NCT05687110). NVB inhibits the ATPase activity of DNA polymerase theta (Polϴ) (Zhou et al. 2021). Inhibition of Polϴ by NVB also been shown to sensitize cells to ferroptosis (Peng et al. 2024), a regulated mode of cell death that can be initiated following oxidative stress, leading to lipid peroxidation and disruption of plasma membrane integrity (Chen et al. 2021). Because AML cells rely on Polϴ to repair DPCs (Vekariya et al. 2023; Chandramouly et al. 2021), we hypothesized that inhibition of Polϴ with NVB would potentiate APR-246-induced cytotoxicity by increasing DPC burden and enhancing cell death specifically in *TP53*-mutated AML while preserving low toxicity. We show here that this combination is highly synergistic in a panel of AML cell lines and primary AML patient samples and that the synergy involves both p53-dependent and p53-independent mechanisms. These results suggest that the combination of APR-246 with NVB may represent a novel strategy tailored for *TP53* mutant AML patients.

## Results

### Synergistic growth suppression by combination of APR-246 with NVB in MOLM13 cells

We first tested the effect of APR-246 in combination with NVB in a panel of isogenic MOLM13 cell lines including cells carrying common *TP53* variants in AML (R175H, Y220C, M237I, R248Q R273H or R282W) as well as a wildtype (WT) *TP53* cells and cells completely lacking *TP53* (null) (Boettcher et al. 2019). We treated cells with increasing concentrations of each compound as monotherapy (APR-246 5 µM to 75µM, NVB 25µM to 150µM) or in combination with concomitant addition of the two agents. Cell growth and viability was assessed after 72 hrs by Trypan blue staining. The sensitivity to APR-246 as monotherapy was significantly different among the eight cell lines (Extra-Sum-of-Squares F-test (F_(21, 104)_ = 16.55, p < 0.0001). As shown in Fig. 1A, this revealed statistically significant differences between the IC50 of each of the mutant cell lines (25-27 µM) and wildtype (36 µM), the IC50 of each of the mutant cell lines and null cells (48 µM) and between the IC50 of wildtype and null cells (adj. p-value < 0.0001). There were no statistically significant differences between IC50 values among mutant cell lines. Thus, these findings demonstrate a preferential cell killing effect of APR-246 towards *TP53*-mutated MOLM13 cells, with a specificity of 94.57% as monotherapy. Treatment with APR-246 as a single agent at 75 µM resulted in almost complete cell death across all MOLM13 cell lines. NVB monotherapy gave comparable dose-response curves across all cell lines regardless of *TP53* mutational status. As a single agent, NVB produced detectable cytotoxic effects only at concentrations above 100 µM. IC50 values were 150 µM or above (Fig. 1B).

**Figure 1.**
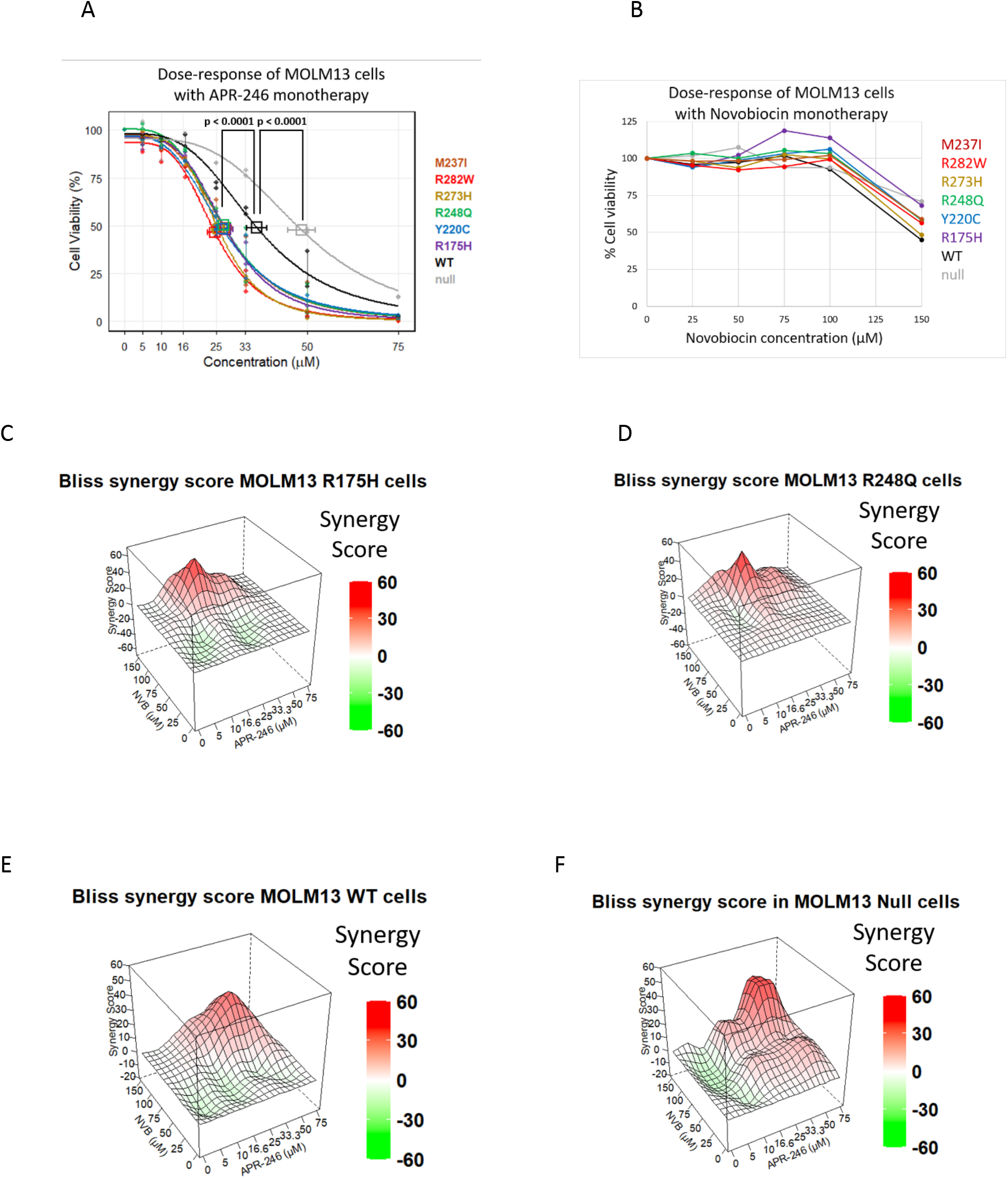
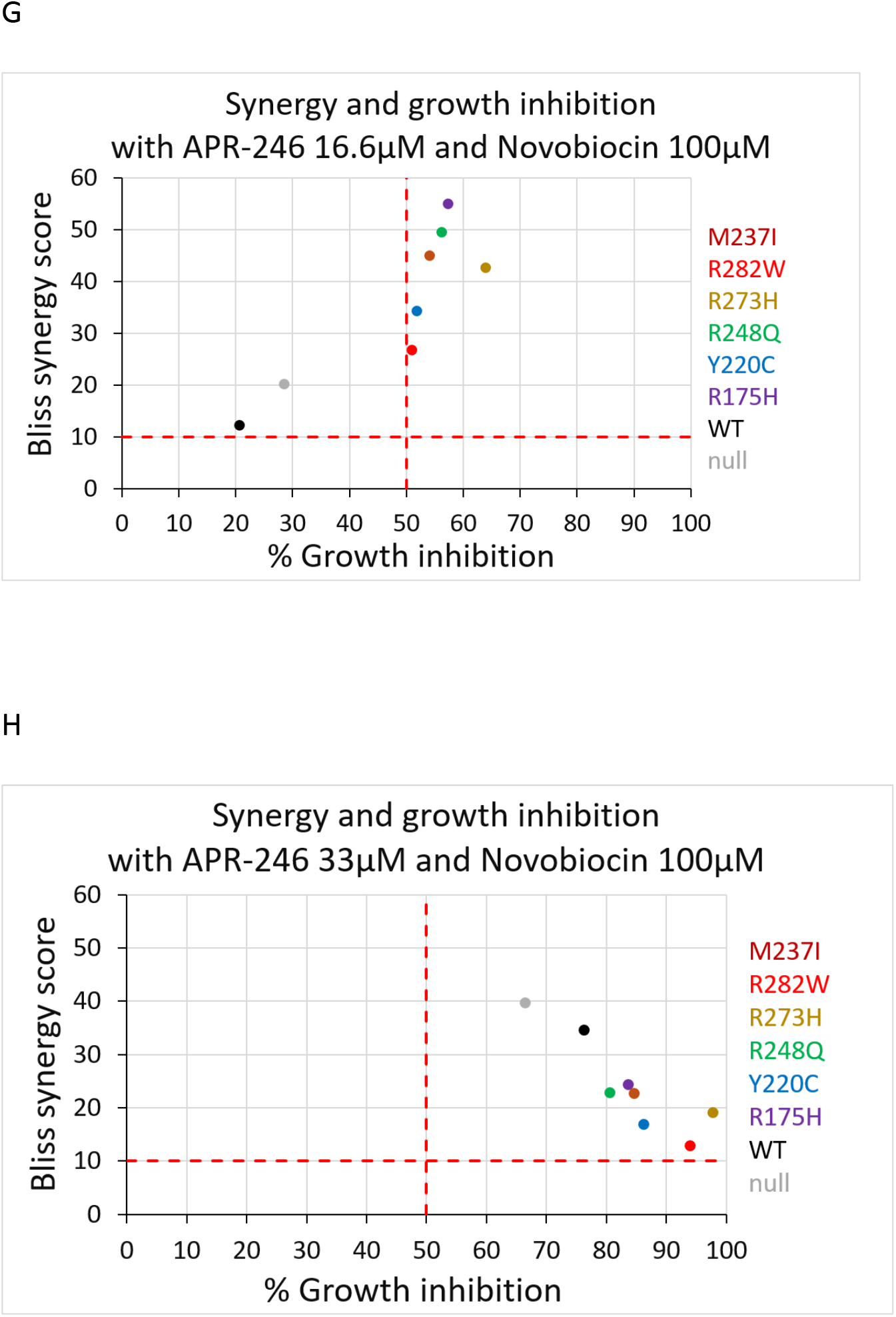
Growth inhibition by APR-246 and NVB in the MOLM13 cell line panel. (A) MOLM13 cells were treated with increasing concentrations of APR-246 and cell viability was assessed after 72 hours by Trypan blue cell count. Data of replicates ws used to generate the dose-response curves with 3-parameter log-logitic model. Dots represent the individual replicates. Squares denote estimated IC50 values and confidence intervals. (B) MOLM13 cells were treated with increasing concentrations of NVB, and cell viability was assessed after 72 hours by Trypan blue cell count. The reported values represent mean of two replicates. (C-F) Bliss synergy scores for the APR-246 and NVB combination treatment. The color scale reflects the type of interaction between the two compounds: antagonistic (green), additive (white), or synergistic (red); a synergy score of 10 or above was considered synergistic; (C) MOLM13 *TP53* R175H cells, (D) MOLM13 *TP53* R248Q cells, (E) MOLM13 *TP53* WT cells, and (F) MOLM13 *TP53* null cells. The reported values represent an average of two independent replicates. (G) Dot plot showing Bliss synergy scores and growth inhibition for all MOLM13 cell lines treated with the combination of APR-246 at 16.6 µM and NVB at 100 µM. (H) Dot blow showing Bliss synergy scores and growth inhibition for all MOLM13 cell lines treated with the combination of APR-246 at 33 µM and NVB at 100 µM.

We observed a strong synergy with the combination of APR-246 and NVB across all cell lines carrying *TP53* missense variants as assessed using the Bliss model. The strongest synergy was observed with the same dose combination of APR-246 and NVB (16.6 µM and 100 µM, respectively). The highest synergy score was recorded in cells carrying *TP53* R175H, followed by R248Q, M237I, R273H, Y220C, and R282W cells (Bliss synergy scores: 55.12, 49.62, 45.06, 42.78, 34.42 and 26.87, respectively; values above 10 indicate synergy) (Fig. 1C, D; Suppl. fig. S1A, S1B; Suppl table S1). The synergy scores of mutant cells did not form a distinct cluster at APR-246 16.6µM with NVB 100µM, and WT and null cells were not identified as having significant outlier synergy scores (p > 0.05, Grubbs’ outlier test).

At the concentrations associated with maximal synergy, growth inhibition was approximately 2-3 fold greater in *TP53*-mutant cells than in *TP53* WT or null cells treated with similar concentrations (p = 0.0248, Wilcoxon Rank Sum test with a continuity correction), while no statistically significant difference was observed between WT and null cells (Fig. 1G; Suppl. fig. S1C, S1D). Importantly, the concentration combination that resulted in maximal synergy among mutant cells was least toxic among WT cells (Fig. 1G), emphasizing the high specificity of this combination.

The combination of the two agents produced synergy also in *TP53* WT and *TP53* null cells (Fig. 1E, F; Suppl. fig. S1A, S1B). However, to achieve maximal synergy, a two-fold higher concentration of APR-246 was required compared to the concentration needed for *TP53* mutant cells, and maximal synergy scores were lower than those observed in the *TP53* mutant cells. The most synergistic combination, i.e. APR-246 at 33.3 µM and NVB at 100 µM, resulted in similar Bliss synergy scores for *TP53* WT and null cells (34.68 and 39.74, respectively; Suppl. table S2). Treatment with APR-246 at 33.3µM and NVB at 100µM caused more potent growth inhibition in *TP53* mutant cells than in WT or null cells. The lower synergy scores in mutant cells reflect the high cell death rates that each compound induces as monotherapy in these concentrations (Fig. 1H; see below).

### NVB induces accumulation of DNA-protein crosslinks

To test if inhibition of Polϴ in the MOLM13 cells would lead to increased DNA-protein crosslinks (DPCs), we assessed formation of DPCs 6 hours after treatment with APR-246 and NVB alone or in combination using the ARK assay (Hu et al. 2020). We compared DPC levels to those induced by formaldehyde in equimolar concentrations to APR-246 (16.6 µM). We observed the highest levels of DPC in cells exposed to formaldehyde, probably because it takes time for APR-246 to be converted to MQ and generate formaldehyde (Lambert et al., 2009), during which some DPCs are being repaired by Polϴ-independent mechanisms (Hu et al., 2020). NVB increased DNA-protein crosslink accumulation in combination with either APR-246 or formaldehyde, consistent with inhibition of PolƟ-mediated DPC repair (Fig. 2A). Of note, treatment with NVB as monotherapy induced DPCs, presumably because MOLM13 which carry FLT3-ITD have inherent high levels of formaldehyde due to aberrant C1 metabolism (Vekariya et al., 2023) (Fig. 2A).

**Figure 2.**
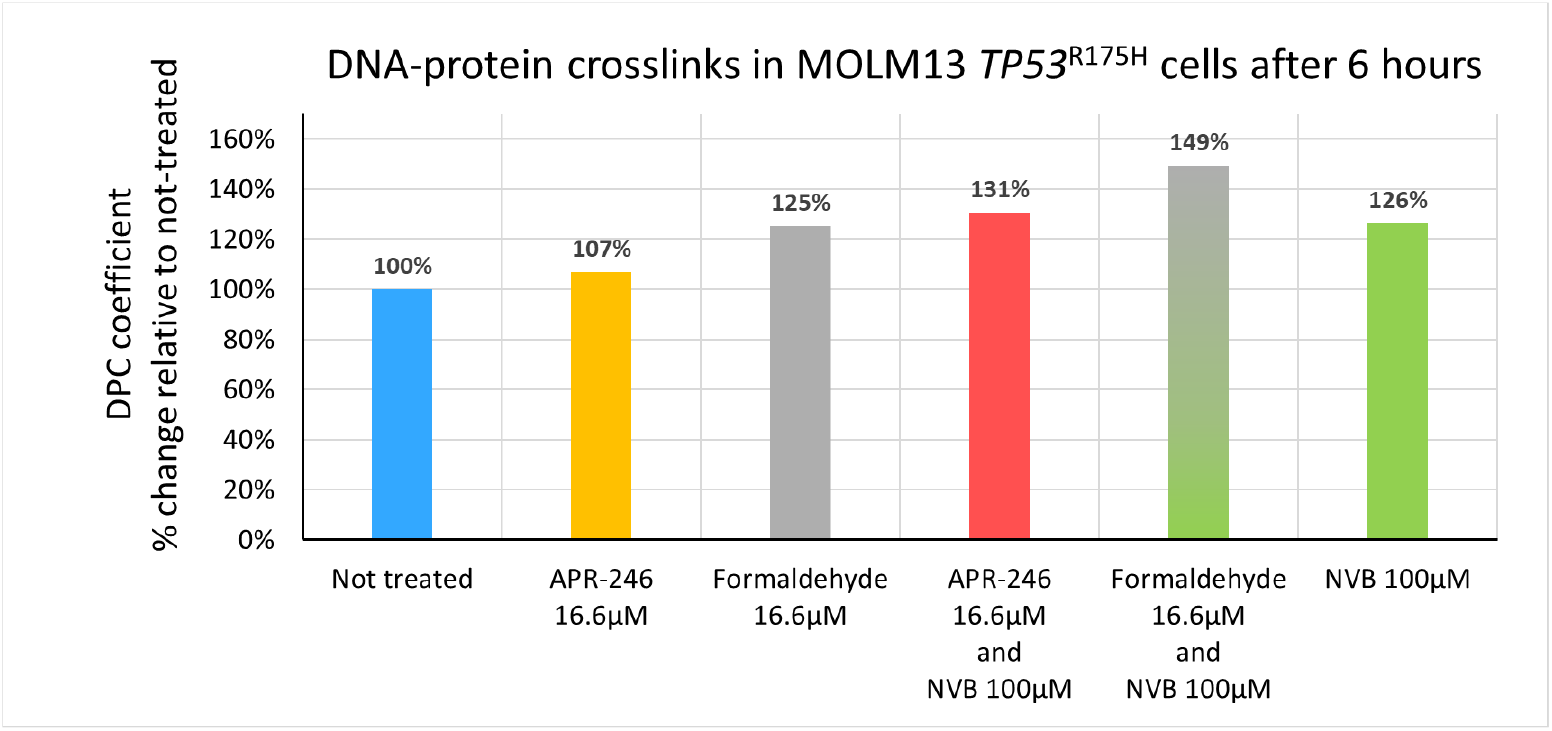
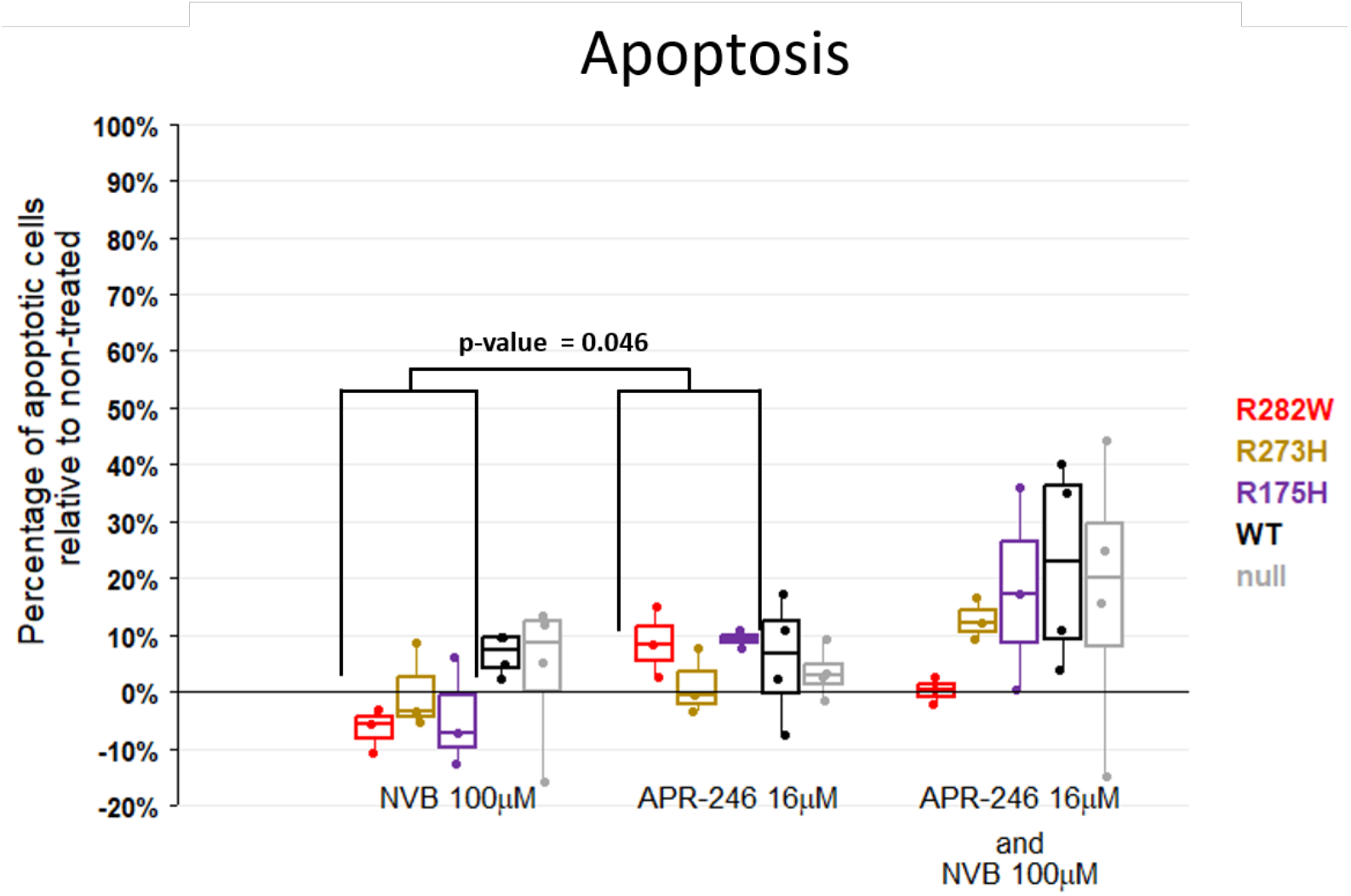
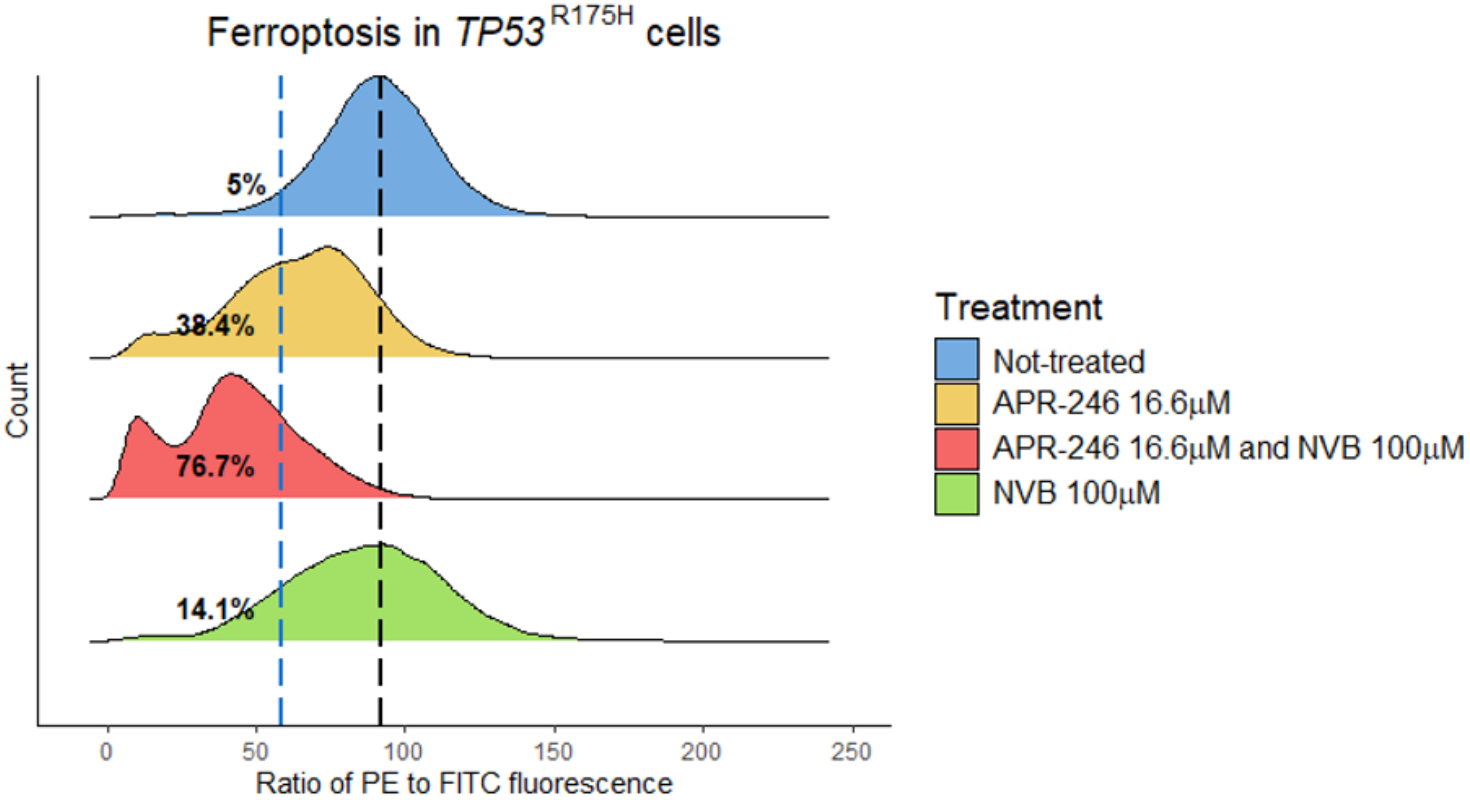
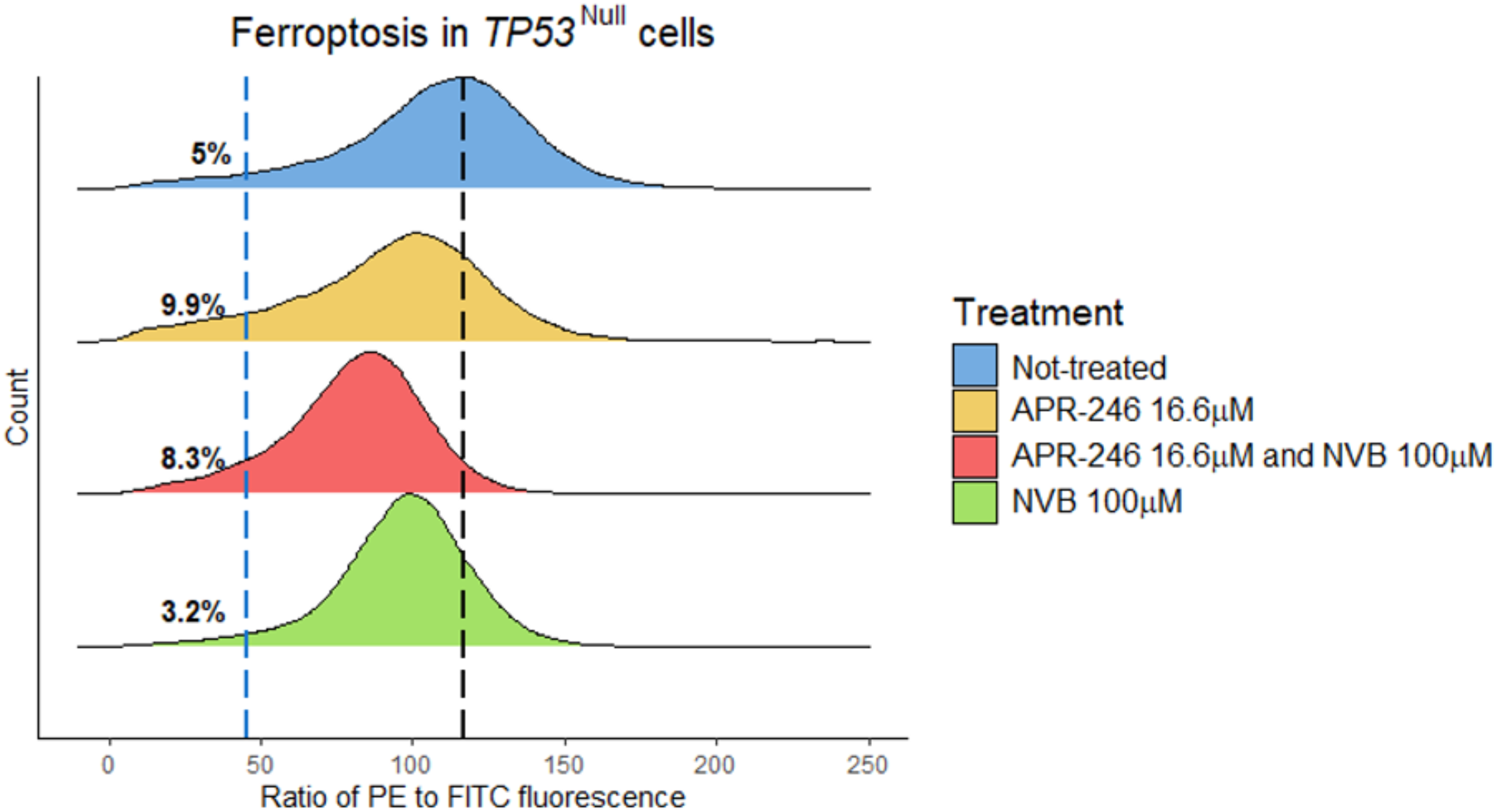
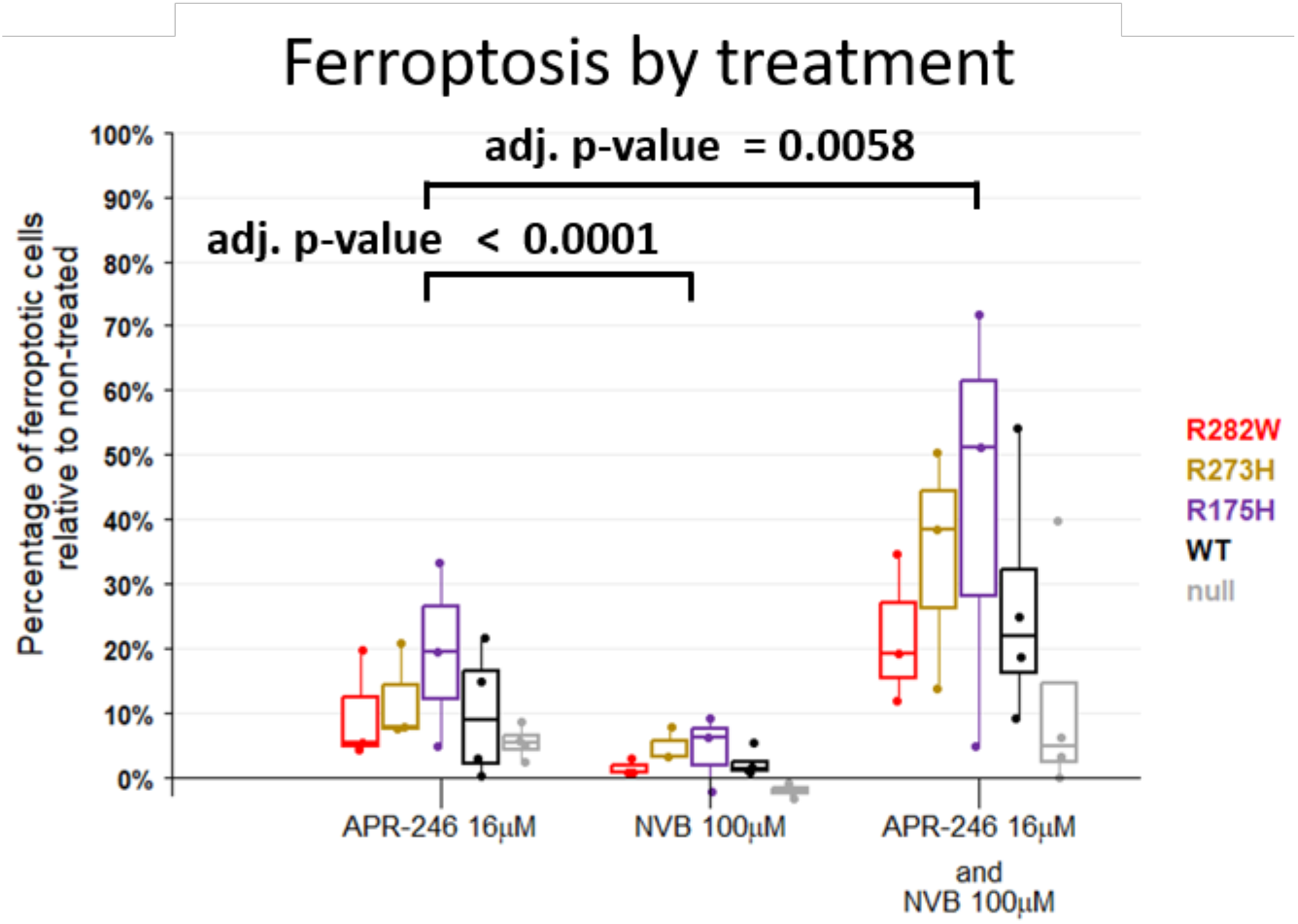
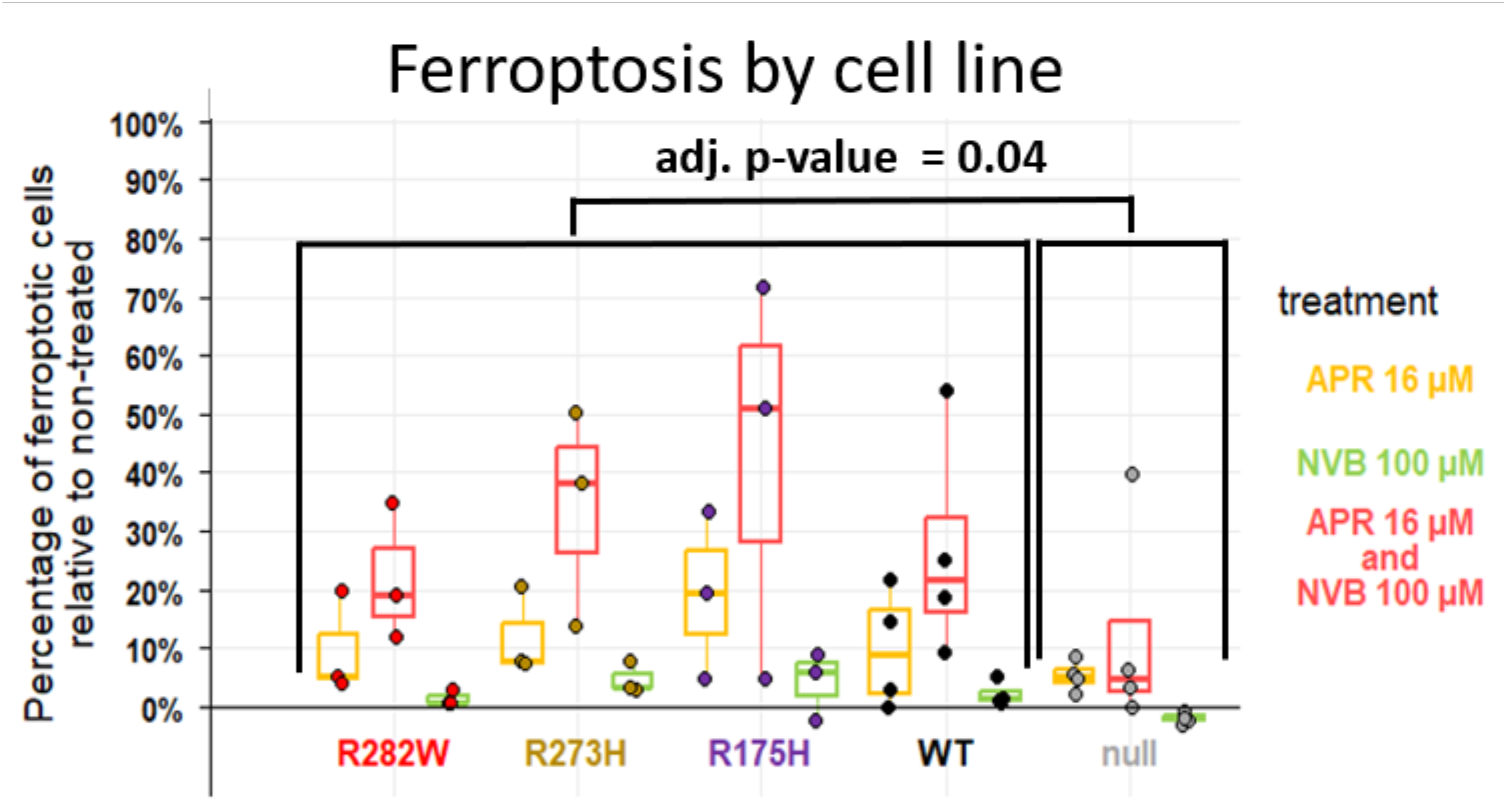
Generation of DNA-protein crosslinks and induction of apoptotic and ferroptotic cell death upon treatment with APR-246 and NVB. (A) Determination of DNA-protein crosslinks (DPCs) in MOLM13 *TP53* R175H cells after 6 hrs of treatment with the indicated agents. Values are percentage change relative to untreated cells. (B) Assessment of apoptosis in MOLM13 *TP53* null, WT, R175H, R273H and R282W cells by Annexin V-FACS. Fraction of Annexin V-positive cells at 24 hours of treatment with the indicated concentrations of APR-246 and NVB alone or in combination is indicated. Each dot represents a biological replicate. Boxes represent the interquartile range (IQR) and medians are shown as bars. Whiskers represent values less than the 25th percentile and greater than the 75th percentiles, and they extend no more than 1.5 × IQR units away from either end of the boxes. p-value of the *post hoc* non-parametric multiple contrast test are presented. (C-D) Two representative replicates of flow cytometry data demonstrating the effect of each treatment over the measured PE to FITC ratio. Lipid peroxidation induces a shift in the emission signal of the BODIPY-C11 reagent from PE to FITC. A lower ratio (shift to the left) represents higher lipid peroxidation. Data that were recorded of 20-30,000 PI-negative cells are shown for each treatment. Non-treated cells (blue) follow a uni-modal distribution (mode is delineated with dashed black line). The PE to FITC value of the 5^th^ percentile of the non-treated cells in each replicate (dashed blue line) was used as a cut-off to quantify the percentage of ferroptotic cells in the treated samples. C – R175H cells, D – null cells. (E-F) Percentage of ferroptotic cells relative to non-treated samples in MOLM13 *TP53* null, WT, R175H, R273H and R282W cells. Each dot represents a biological replicate. Boxes represent the interquartile range (IQR) and medians are shown as bars. Whiskers represent values less than the 25th percentile and greater than the 75th percentiles, and they extend no more than 1.5 × IQR units away from either end of the boxes. Adjusted p-value of the *post hoc* test are presented. E – data stratified by treatment. F – data stratified by cell line.

### Cell death induced by combination treatment with APR-246 and NVB

APR-246 can induce both apoptotic and ferroptotic cell death (Mohell et al. 2015; Birsen et al. 2022; Fujihara et al. 2022). To determine whether the observed growth inhibition was a consequence of induction of apoptosis or ferroptosis or both, we assessed Annexin V staining (apoptosis; Suppl. fig. S2) and the PE to FITC shift in emission of BODIPY-C11 reagent (where a lower PE to FITC ratio reflects increase in lipid peroxidation) concomitantly by flow cytometry 24 hours after exposure to APR-246 and NVB at the most synergistic concentrations (16.6 µM and 100 µM, respectively), both as monotherapy and in combination, and focused on five cell lines: R175H (that demonstrated highest synergy), R273H (that demonstrated the highest sensitivity), R282W (that demonstrated the lowest sensitivity among mutant cells), WT cells and null cells.

APR 16.6µM monotherapy induced a significant increase in apoptosis by up to 8% in the *TP53*-mutant cells as compared to NVB 100µM monotherapy (relative effect = 0.864, 95% CI: 0.510 to 1.218, p-value = 0.046). The combination of APR-246 and NVB caused increased apoptosis by up to 20% in all cells except R282W cells as compared to APR monotherapy (3.9-fold increase from a median of 4.8% to 18.7%) but this did not reach statistical significance (Fig. 2B). WT and null cells showed similar levels of apoptosis when comparing NVB monotherapy to APR monotherapy (p-value = 0.9). R282W cells deviated from the main pattern with reduced percentage of apoptotic cells when NVB was added to APR-246 16.6µM (relative effect = 0.838, 95% CI: 0.583 to 1.094, p-value = 0.019). The difference between the WT and null cells on the one hand and the mutant cells on the other hand did not reach statistical significance (p-value = 0.26). These data demonstrate several important points. First, NVB as monotherapy induces low levels of apoptosis in both WT and null cells, highlighting a weak p53-independent mechanism. Second, mutant cells are resistant to this effect (as also shown in the dose-response curves for NVB monotherapy; Fig. 1B), probably due to their ability to withstand the DNA damage induced by NVB. Third, APR-246 at 16.6µM induces low levels of apoptosis in all cells, but levels are slightly higher in mutant cells (approx. in 8.7% in R175H and R282W cells) than in WT or null cells (6.6% and 2.9%, respectively). Fourth, combination of APR-246 and NVB led to a non-significant increase in apoptosis in all cells, and most prominently in WT and null cells (23% and 20%, respectively), emphasizing the p53-independence of this effect.

To quantify the percentage of ferroptotic cells, we used as a cut-off the PE to FITC ratio value of the 5^th^ percentile in the non-treated sample of that replicate (Fig. 2C, D). All three treatments were statistically distinct in their ability to induce ferroptosis (Fig. 2E). NVB 100µM as monotherapy induced minimal levels of ferroptosis above base line untreated levels (raw mean 2.17 ± 1.2% Standard Error of the Mean, SEM), whereas APR-246 at 16.6µM as monotherapy induced ferroptosis in 11.2 ± 2.27% of the cells (a 5.17 fold increase when compared with NVB monotherapy, adj. p-value < 0.0001, Cohen’s *d* = 1.7). The combination of APR-246 16.6 µM and NVB 100µM induced ferroptosis in 27.56 ± 5.17% of the cells (a 2.45 fold increase when compared with APR-246 monotherapy, adj. p-value= 0.0058, Cohen’s *d* = 1.17) (Suppl. table S3A). Levels of ferroptosis were 3.18-fold higher in the pooled group of mutant cells when compared with null cells (adj. p-value = 0.044, Cohen’s *d* = 1.05), and this was mainly due to higher levels of ferroptosis among R175H cells (4.2-fold) and R273H cells (3.24-fold) when compared with null cells (Cohen’s *d* = 1.5 and 1.12, respectively, adj. p-value = 0.044 for each comparison). Interestingly, WT cells had 2.47-fold higher levels of ferroptosis when compared with null cells (adj. p-value = 0.044, Cohen’s *d* = 0.97), but these levels were not statistically different from ferroptosis levels in mutant cells, implying that, collectively, p53-expressing cells are more prone to ferroptosis. This was confirmed by statistically significant 3-fold higher levels of ferroptosis among p53-expressing cells when compared with null cells (adj. p-value = 0.044, Cohen’s *d* = 1.03) (Fig. 2F; Suppl. table S3B).

Thus, combination of NVB and APR-246 leads to a significantly higher levels of ferroptosis with a 2.45-fold increase as compared to APR-246 monotherapy and a 12.6-fold increase as compared to NVB monotherapy (Suppl. table S3A) regardless of *TP53* mutational status. Furthermore, 3-fold higher levels of ferroptosis were found in p53-expressing cells, regardless of treatment, pointing to a p53-dependent predisposition to ferroptosis (Suppl. table S3B). The latter effect is presumably driven mainly by differences between R175H and R273H cells and null cells and correlates with the levels of synergy in these cells. In summary, our results demonstrate that the combination of APR-246 and NVB induces both apoptosis and ferroptosis in the MOLM13 cells, and that *TP53*-mutant and WT cells are more prone to ferroptosis.

### APR-246 and NVB synergize to inhibit growth of primary CMML/AML patient samples

Having demonstrated potent combinatorial synergy between APR-246 and NVB in AML cell lines, we next assessed potential synergy in primary patient samples with different *TP53* status. We evaluated peripheral blood or bone marrow samples from six patients, including three *TP53*-wild-type CMML patients and three *TP53*-mutated AML/CMML patients (Table 1). We tested viability using the CellGlo ATP-based assay three days after incubation with APR-246 (0-75µM) and NVB (0, 50µM or 100µM). *TP53*-mutated samples demonstrated substantially greater sensitivity to APR-246 monotherapy than *TP53*-wild-type samples, with IC50 values below 2 µM in all mutant samples compared with more than 15 µM in wild type samples (Table 1). Notably, two *TP53*-mutated samples demonstrated nanomolar IC50 values (14 nM and 222 nM), while in two TP53-wild-type samples the IC50 was not reached even with 75µM of APR-246 (Fig. 3A, B).

**Figure 3.**
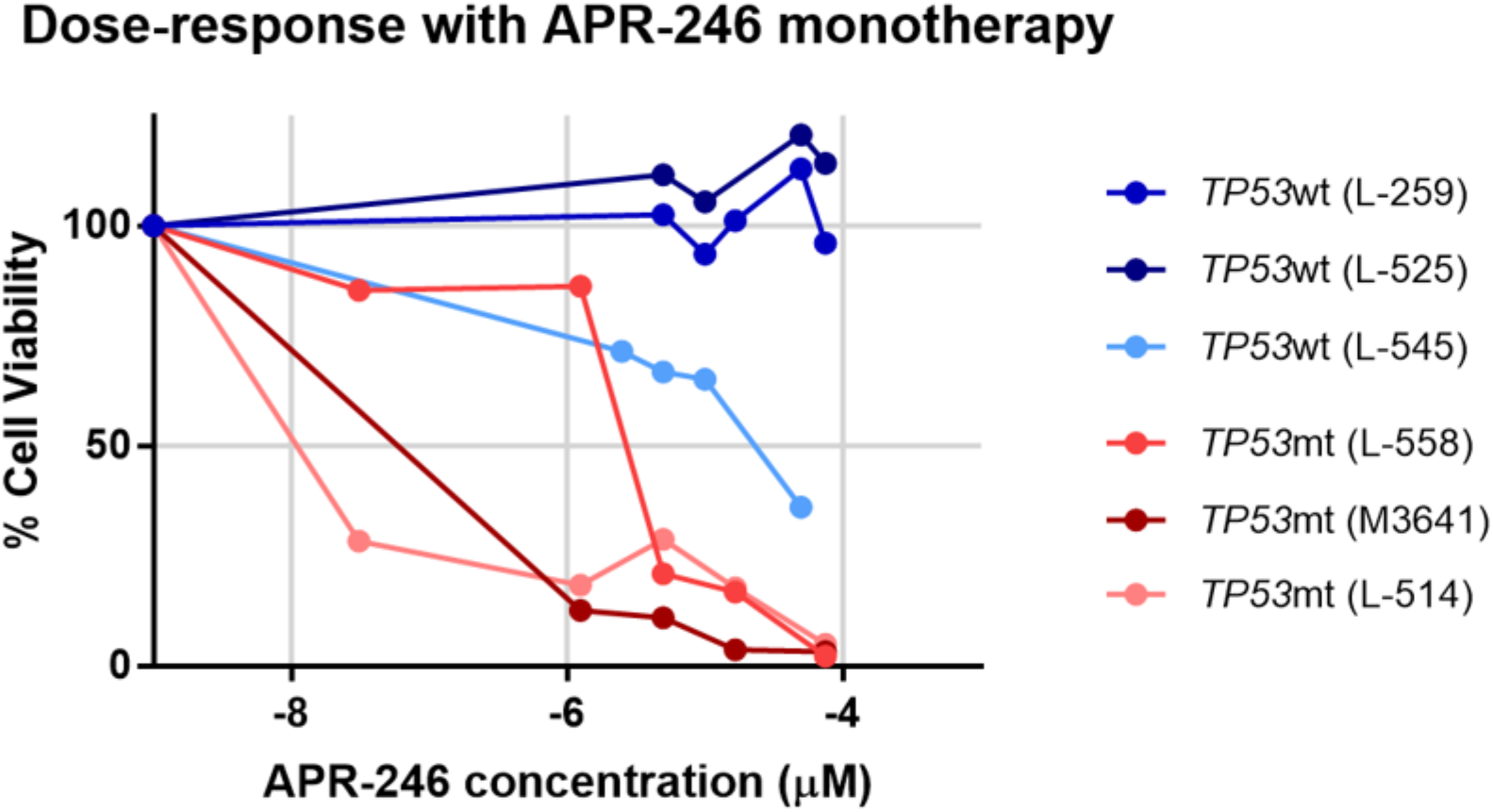
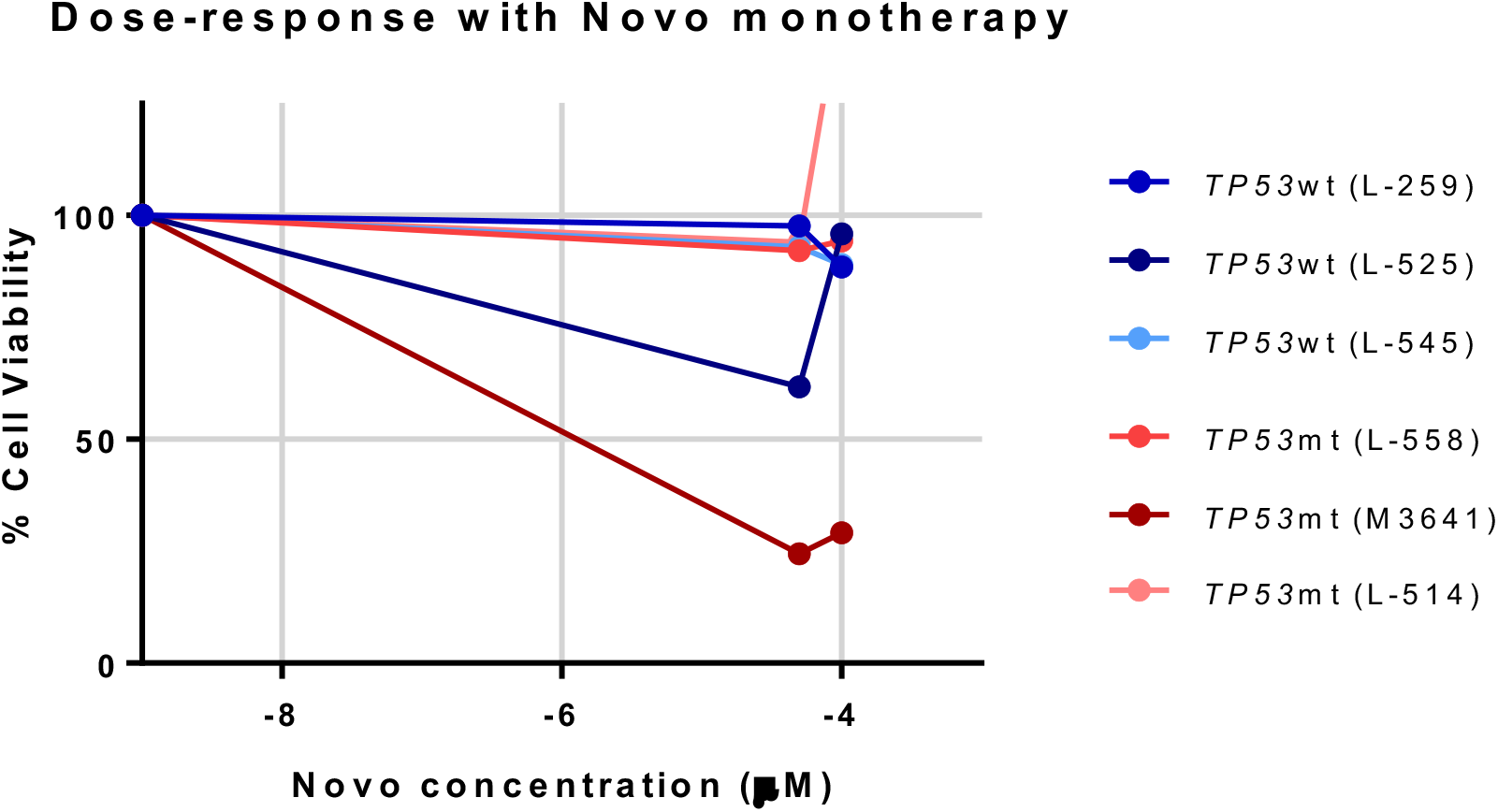
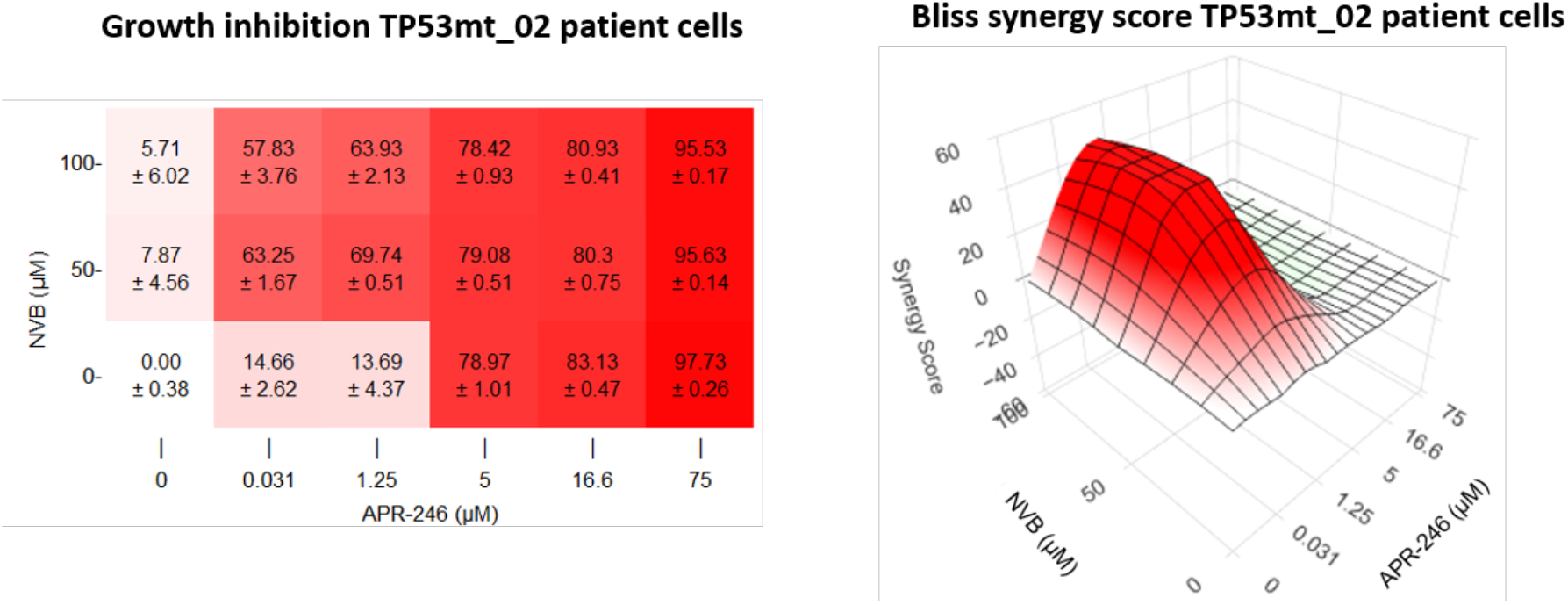
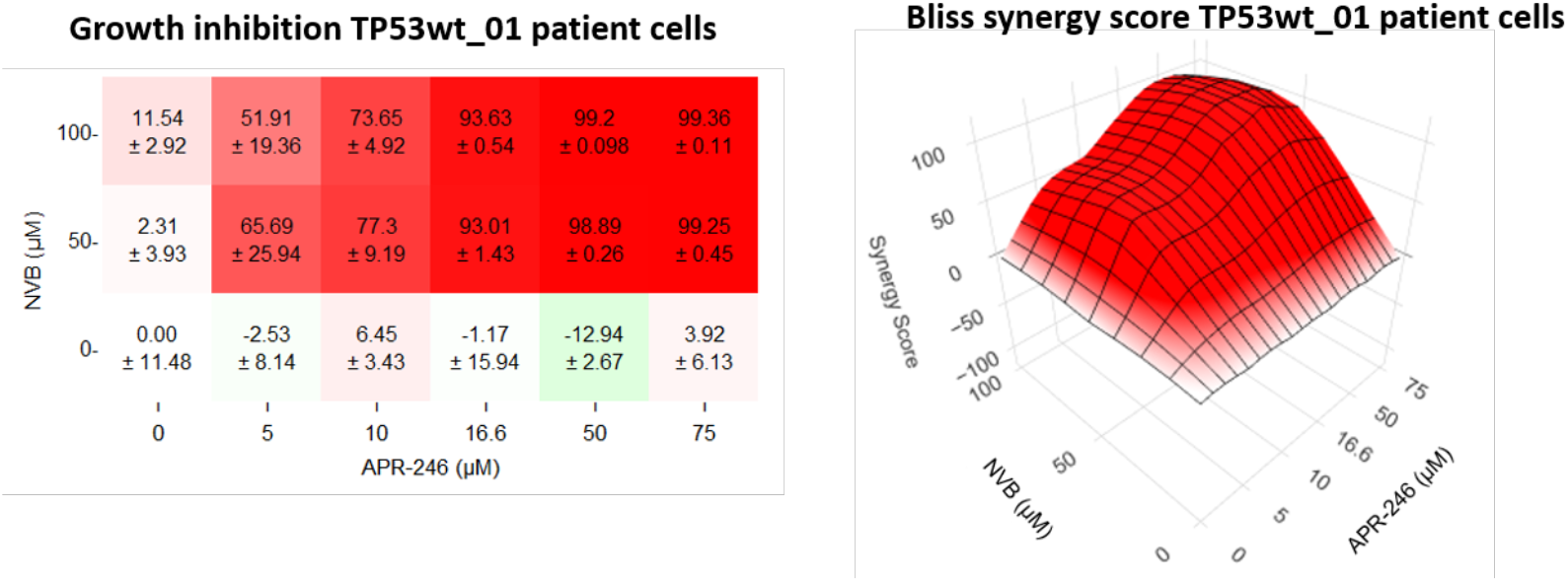
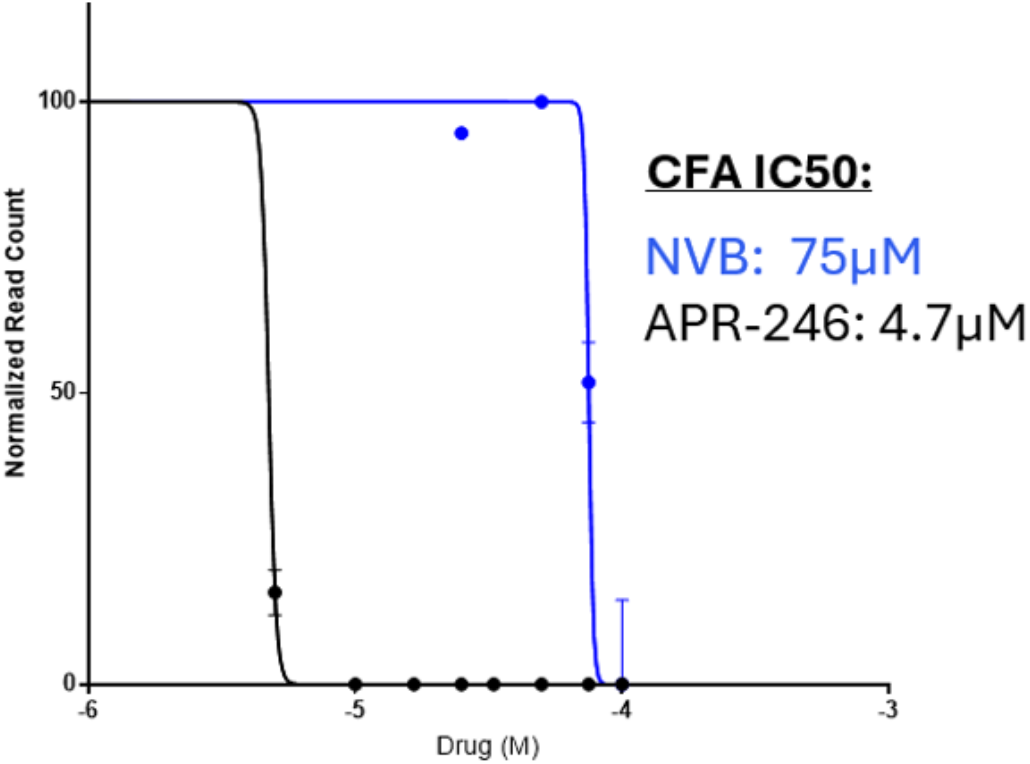
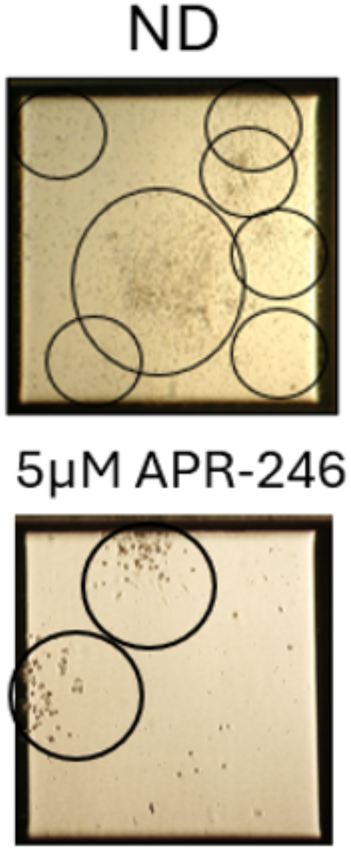
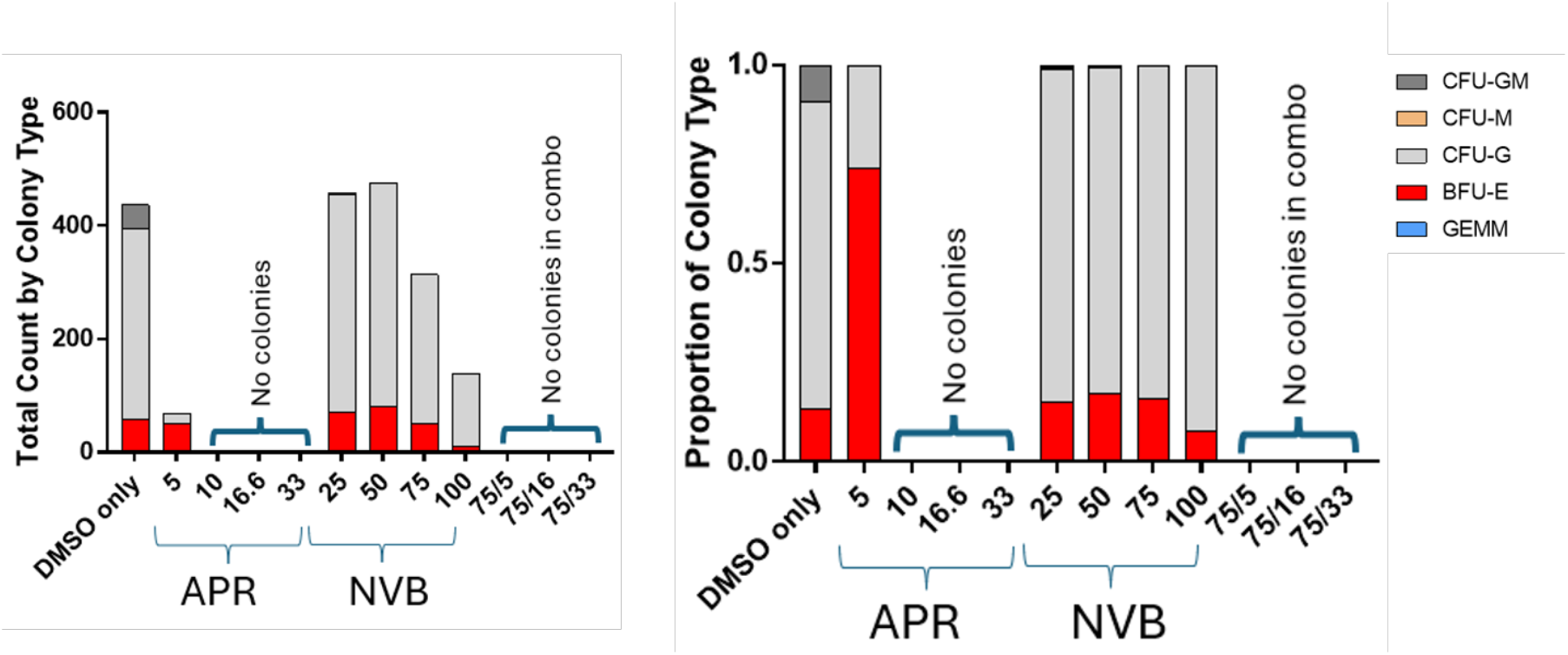
Growth suppression of primary AML patient samples by combination treatment with APR-246 and NVB. Dose response curves of APR-246 (A) and NVB (B) as monotherapies. ATP-based viability assay triplicate average values showing normalized (to non-treated cells) percentage viability. (C) A representative *TP53*-mutated primary sample. Left panel: Heatmap of normalized percentages of growth inhibition. Right panel: Bliss synergy scores for each of the tested dose combinations. (D) Same as in C, for a representative *TP53*-wild type primary sample. Synergy is observed in both samples with higher efficacy (lower drug concentrations) in the *TP53*-mutated sample. Panels (E-G): Bone marrow mononuclear cells were plated for colony forming assay (CFA) on one TP53mt case (therapy related high risk MDS carrying: *TP53* c.404G>A, C135Y (41%); *TP53* c.422G>A,C141Y (3%); *TP53* c.530C>G, P177R (45%) with complex karyotype in the presence of APR-246 and Novobiocin. (E) IC50 for each single agent. (F) Representative colony photos. (G) Total count of colony by type (left panel) and proportion of colony type present by concentration (right panel). Drug combinations include NVB at 75 µM and APR-246 at 5, 16.6, and 33 µM (75/5, 75/16, and 75/33). Colony types: BFU-E = erythroid, CFU-GM = granulocyte/monocyte mixed, CFU-G = granulocyte, CFU-M = monocyte/macrophage, GEMM = primitive mixed erythroid/myeloid). ND = no drug.

**Table 1.** Primary patient sample information. Clinical characteristics, mutational profile, IC50 for APR-246 as monotherapy, maximal Bliss synergy score and the drug combinations that resulted in maximal synergy.

|  | **TP53wt_01** | **TP53wt_02** | **TP53wt_03** | **TP53mt_01** | **TP53mt_02** | **TP53mt_03** |
|---|---|---|---|---|---|---|
| **Age** | 75 | 72 | 87 | 67 | 85 | 24 |
| **Sex** | Male | Male | Male | Female | Female | Female |
| **WHO dx** | CMML-0 | CMML-1 | CMML-1 | s-AML | CMML-1 | AML |
| **Hb g/L** | 8 | 9.3 | 13.1 | 7.9 | 14.1 | 9.9 |
| **Hematocrit (%)** | 25.6 | 27.5 | 42 | 23 | 48.4 | 30.2 |
| **WBC x 10<sup>9</sup>/L** | 5.7 | 15.3 | 3.4 | 0.5 | 9.8 | 46.5 |
| **Platelet x 10<sup>9</sup>/L** | 170 | 177 | 103 | 18 | 222 | 90 |
| **PB Blasts (%)** | 0 | 0 | 0 | 0 | - | - |
| **BM blasts (%)** | 2 | 5 | 1 | 96 | - | - |
| **Cytogenetics** | 46,XY[20] | 46,XY[20] | 46,XY[20] | 45-46,X,del(X)(q11), add(9)(q22), del(17)(p11.2),-20,+0-1r[cp17]/46,XX[3] | 46,XX[20] | +3,+4 add(5) in 11/20 metaphases |
| **TP53 mutation(s) (VAF)** | None detected | None detected | None detected | ***TP53*** c.742C>T; R248W (84%) | ***TP53*** c.818G>A; R273H (22%) | ***TP53*** c.742C>T; R248W (51%)<br>***TP53*** c.743G>A; R248Q (9%)<br>***TP53*** c.743G>T; R248L (6%)<br>***TP53*** c.775G>T; D259Y (9%) |
| **Other NGS mutations (VAF)** | *TET2* N275Ifs\*18 (42%)<br>*TET2* A1882Efs\*5 (41%) | *EZH2* c.1851+2T>C; p.? (13%)<br>*TET2* K1299N (99%)<br>*ZRSR2* K26\* (66%)<br>*ZRSR2* c.203+1G>A; p.? (9%)<br>*ZRSR2* c.558-2A>C; p.? (10%) | *ASXL1* G646Wfs\*12 (15%)<br>*TET2* Q278\* (28%)<br>*TET2* K751Ifs\*61 (17%)<br>*TET2* Q947\* (18%)<br>*CBL* c.1096-7A>G; p.? (3%)<br>*SRSF2* P95H (50%) | *DNMT3A* R882H (47%)<br>*ETNK1* N244S (45%)<br>*KRAS* A146V (41%)<br>*U2AF1* S34F (45%)<br>*RUNX1* R204L (44%)<br>*STAG2* c.2534-1G>A;p.? (46%)<br>*TET2* D1314Mfs\*49 (48%) | *JAK2* V617F (35%)<br>*KRAS* G12S (16%)<br>*CBL* L380P (3%)<br>*PPM1D* L474Afs\*5 (3%)<br>*TET2* Q591Sfs\*10 (44%) |  |
| **APR-246 Monotherapy IC<sub>50</sub>** | not met | not met | 15.4µM | 14nM | 1.82µM | 222nM |
| **Maximal Bliss Score** | Max. Bliss score: 108.9<br>APR-246: 50µM<br>NVB: 50µM | Max. Bliss score: 327.8<br>APR-246: 16.6µM<br>NVB: 100µM | Max. Bliss score: 3.773<br>APR-246: 50µM<br>NVB: 100µM | Max. Bliss score: 29.12<br>APR-246: 0.031µM<br>NVB: 100µM | Max. Bliss score: 49.17<br>APR-246: 1.25µM<br>NVB: 50µM | no synergy |

Consistent with observations in MOLM13 cells, APR-246 and NVB demonstrated synergistic activity in both *TP53*-mutated and *TP53*-wild type samples; however, synergy occurred at markedly lower APR-246 concentrations in *TP53*-mutant samples (0.031-1.25 µM versus 16.6-50 µM in *TP53*-wild-type samples). Importantly, the combination achieved 70-90% growth inhibition at APR-246 concentrations as low as 1.25-5µM (Table 1, Fig. 3C, Suppl. fig. S3A). In the “TP53mt_01” sample, APR-246 monotherapy was highly potent (IC50 = 14nM), and the highest synergy score was observed with APR-246 0.031µM and NVB 100µM. This likely reflects both the potent single-agent activity of APR-246 and the unexpected proliferative effect observed with NVB 100µM alone (150% viability relative to the non-treated cells, Suppl. fig. S3A), which amplified the calculated synergy score. Notably, APR-246 31nM combined with NVB 50µM resulted in near-complete elimination of viable cells (95% growth inhibition; Supplementary Fig. S3A). Because APR-246 alone already produced near-maximal inhibition at very low concentrations, the ability of the synergy algorithm to identify additional combinatorial effects was likely limited. *TP53*-mutated sample 3 (“TP53mt_03”) did not demonstrate measurable synergy, likely because APR-246 monotherapy already induced 91% growth inhibition at 2.5µM (Suppl. fig. S3A), leaving little opportunity for additional improvement with combination treatment.

Interestingly, wild type samples demonstrated highly heterogeneous synergy scores. Two samples (“TP53wt_01” and “TP53wt_02”) had very high synergy scores of 109 and 328, respectively. These high synergy scores likely reflect the limited activity of either agent as monotherapy, resulting in large combination effects relative to the expected additive response (IC50 not reached either for APR-246 or for NVB). In sample “TP53wt_03” the combination of these two compounds demonstrated no synergy (maximal synergy score of less than 10) and induced 64% growth inhibition with high concentrations of APR-246 (50µM) which were only slightly increased to 72% with the addition of NVB 100µM (Table 1, Fig. 3D, Suppl. fig. S3B).

Collectively, these findings demonstrate that APR-246 and NVB exhibit synergistic activity in primary myeloid samples, with substantially greater potency observed in *TP53*-mutated cells. To evaluate the effect of this combination on clonogenic potential, we performed colony-forming assays using bone marrow mononuclear cells from a therapy-related high risk MDS patient harboring bi-allelic *TP53* alterations and a complex karyotype. The IC50 values for APR-246 and NVB monotherapy were approximately 5 µM and 75 µM, respectively (Fig. 3E). Note, we did not test APR-246 concentrations lower than 5µM in combination with NVB, but all colonies were eliminated when NVB 75µM was added to 5µM of APR-246, again underscoring the enhanced efficacy of this therapeutic approach (Fig. 3F, G).

### Cell priming with NVB enhances both synergy and growth inhibition but requires a higher APR-246 dose

Since APR-246 is converted to MQ during the first 24 hours of its administration (Lambert et al. 2009), we hypothesized that both DNA damage produced by APR-246-generated formaldehyde and oxidative stress could be increased if Polϴ was already inhibited by the time of APR-246 administration. Therefore, we also tested a different experimental set-up in which MOLM13 *TP53* R175H cells were treated with NVB 24 hours prior to the addition of APR-246 and assessed viability 72 hours after APR-246 administration (Fig. 4A). To maintain the cells in log-phase during the entire extended experimental period, we reduced seeding density to 200,000 cells/mL. During the first 24 hours cells proliferated, such that at the time of APR-246 administration cells reached 350,000 cells/mL. The cell killing effect of APR-246 is influenced not only by its concentration but also by the cell density at the time of treatment; the efficacy of APR-246 decreases at higher cell density, as illustrated in Fig. 4B. We therefore used as control cells that were seeded at similar density (200,000 cells/mL) and exposed to the same concentrations of the two drugs that were administered concomitantly 24 hours after seeding (Fig. 4A). This allowed a direct comparison of the two treatment protocols, as results were measured at the same time point (72 hrs) after the addition of APR-246. As shown in Fig. 4C and D, the sequential treatment achieved similar growth inhibition with APR-246 16.6 µM and NVB 100 µM as the concomitant treatment (47.74% and 48.8%, respectively). However, APR-246 at 33.3 µM and NVB at 100 µM elicited a nearly complete growth inhibition (93.16%) (Fig. 4C), as compared to 66.74% inhibition when administered concomitantly (Fig. 4D). The maximal Bliss synergy for the sequential treatment regimen achieved by APR-246 33.3 µM and NVB 100 µM was nearly two-fold greater (45.11) (Fig. 4E, F) than the maximal score obtained for the concomitant treatment with lower concentrations of APR-246 16.6 µM and NVB 100 µM (27.02) (Fig. 4G, H). The dot plot summarizes the higher potency of the APR-246-NVB sequential treatment with regard to both Bliss synery and growth inhibition (Fig. 4I).

**Figure 4.**
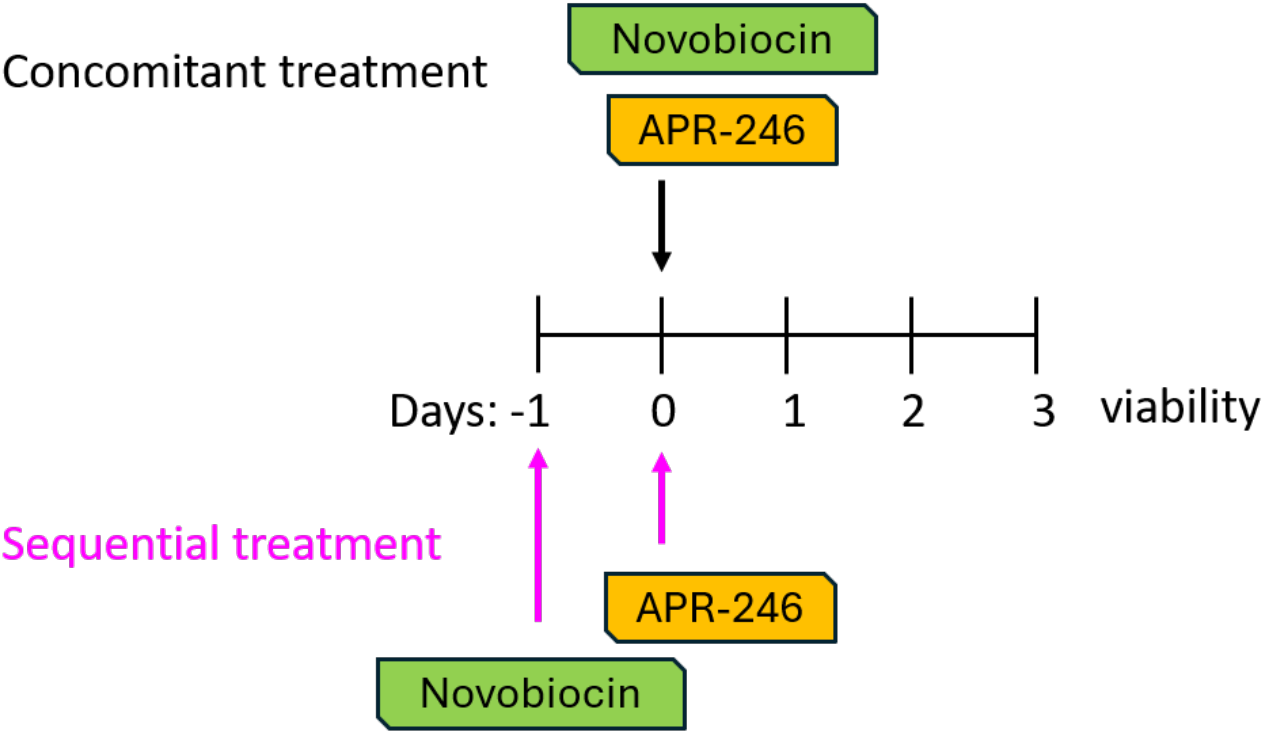
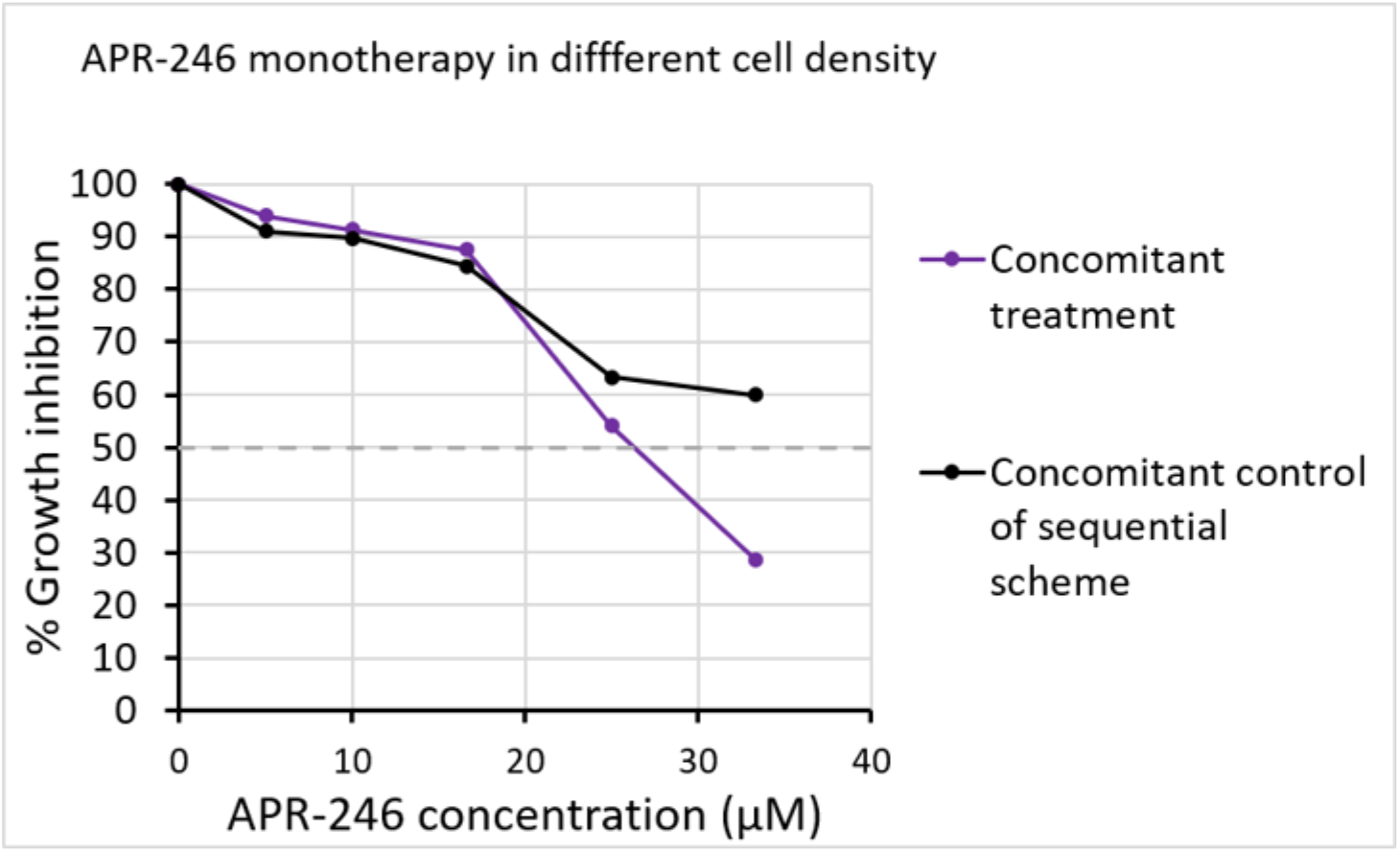
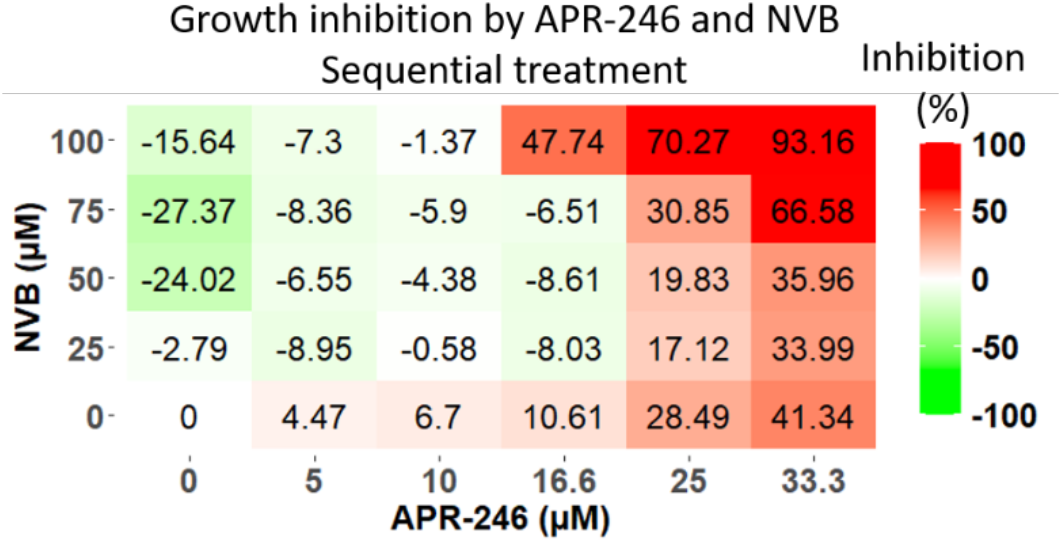
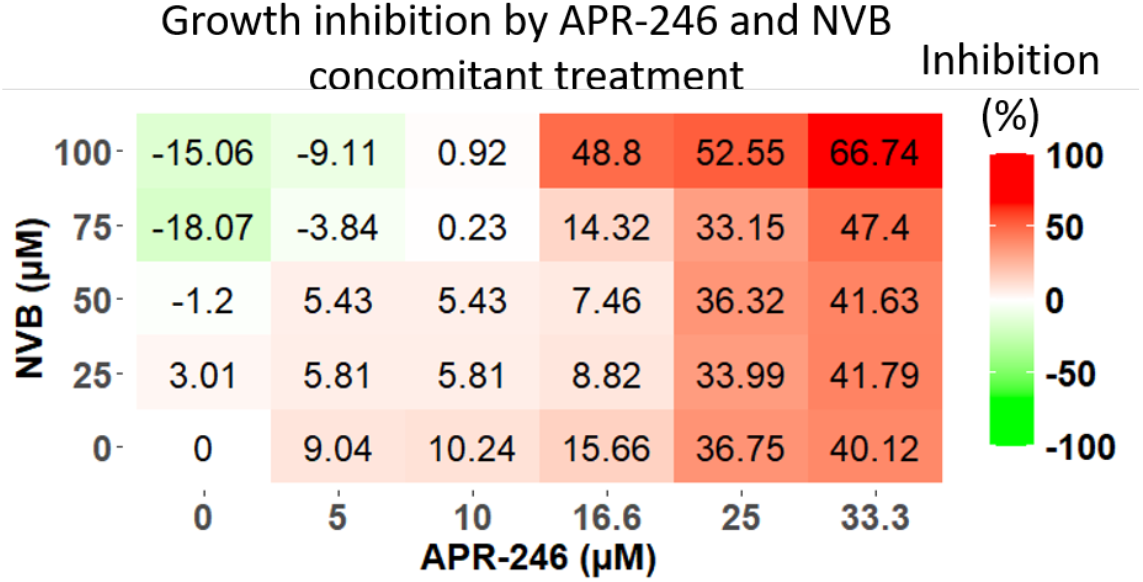
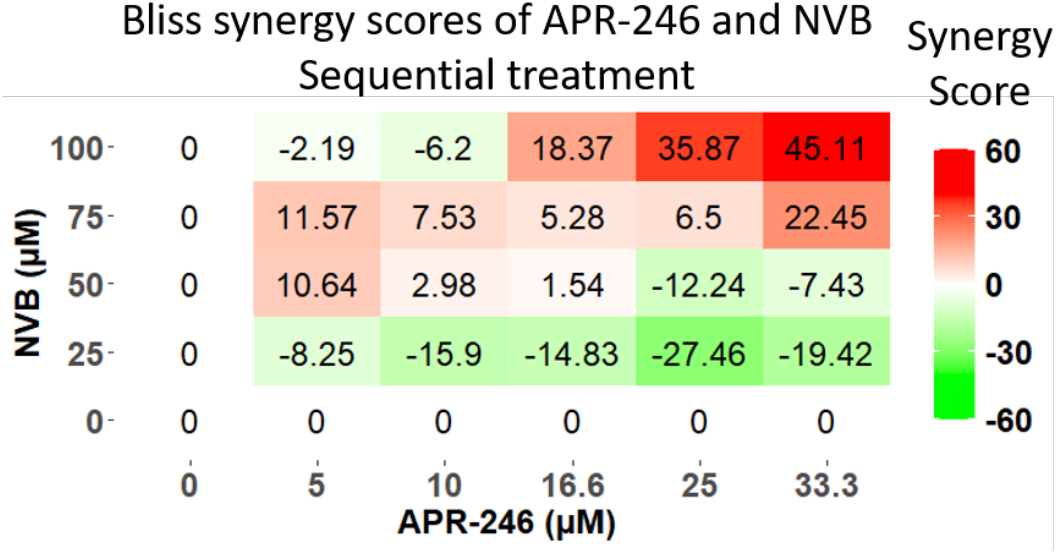
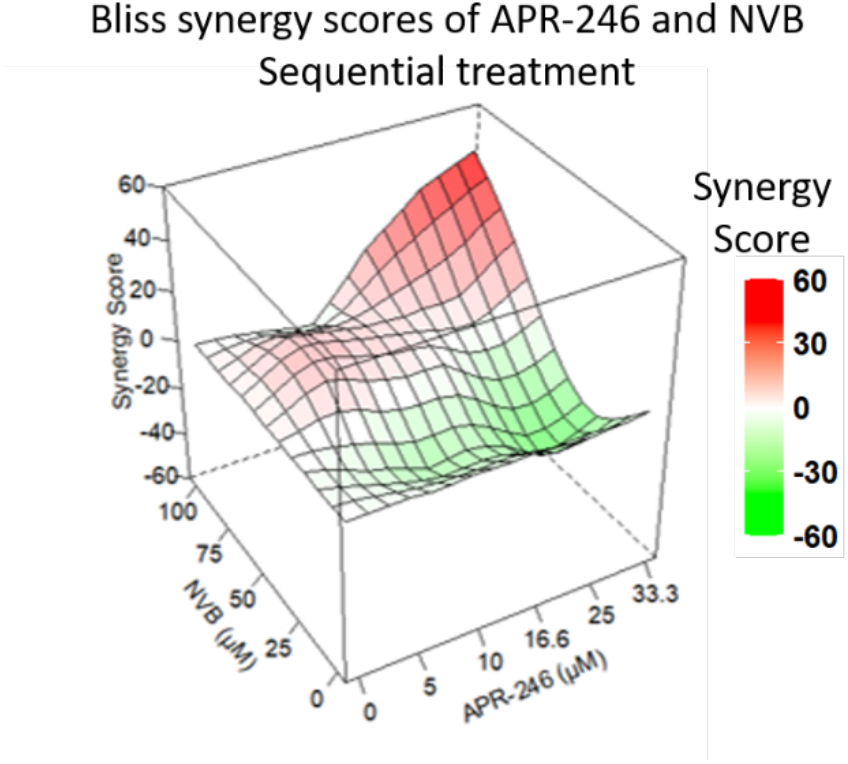
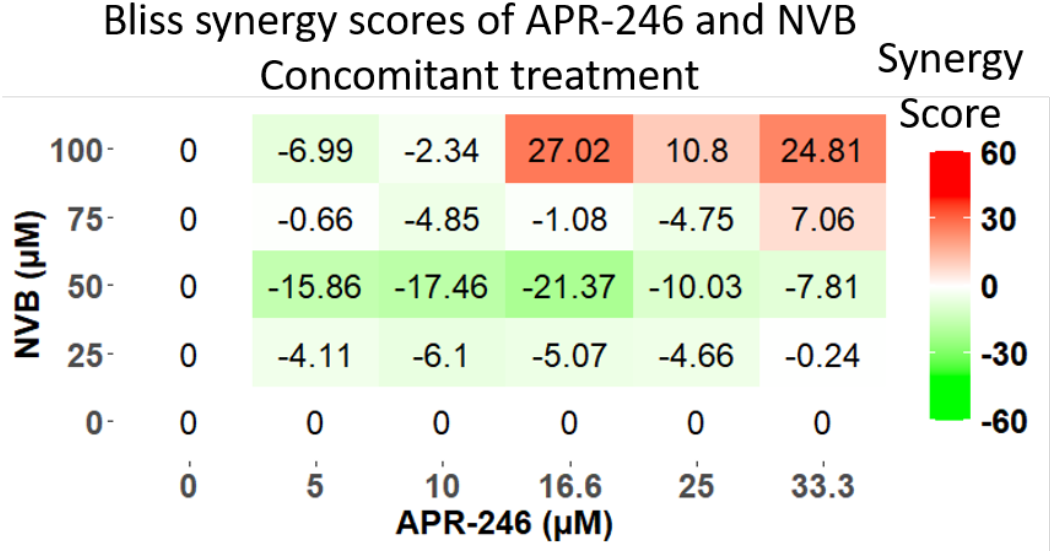
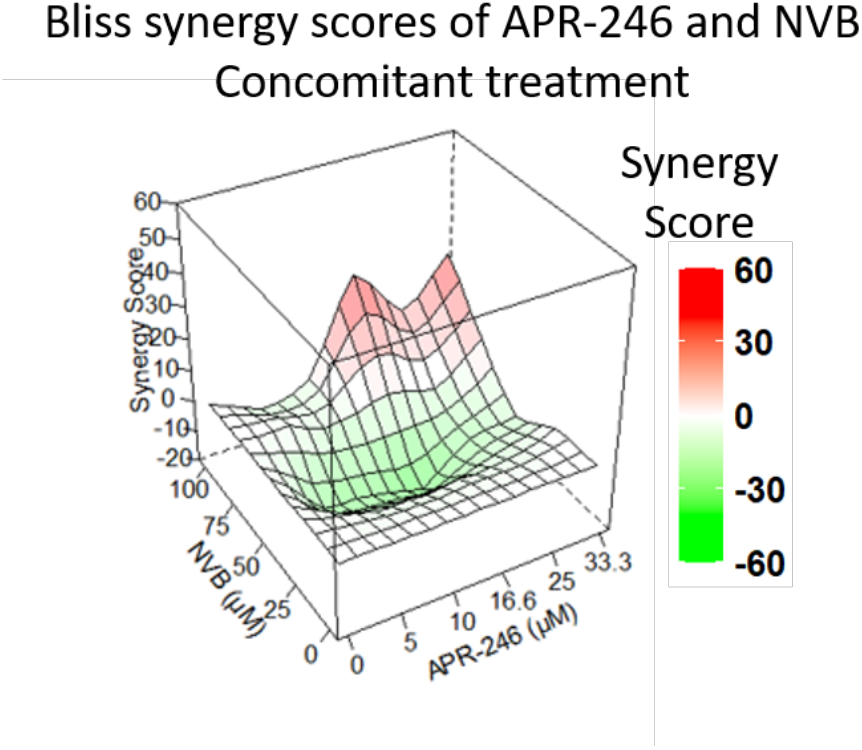
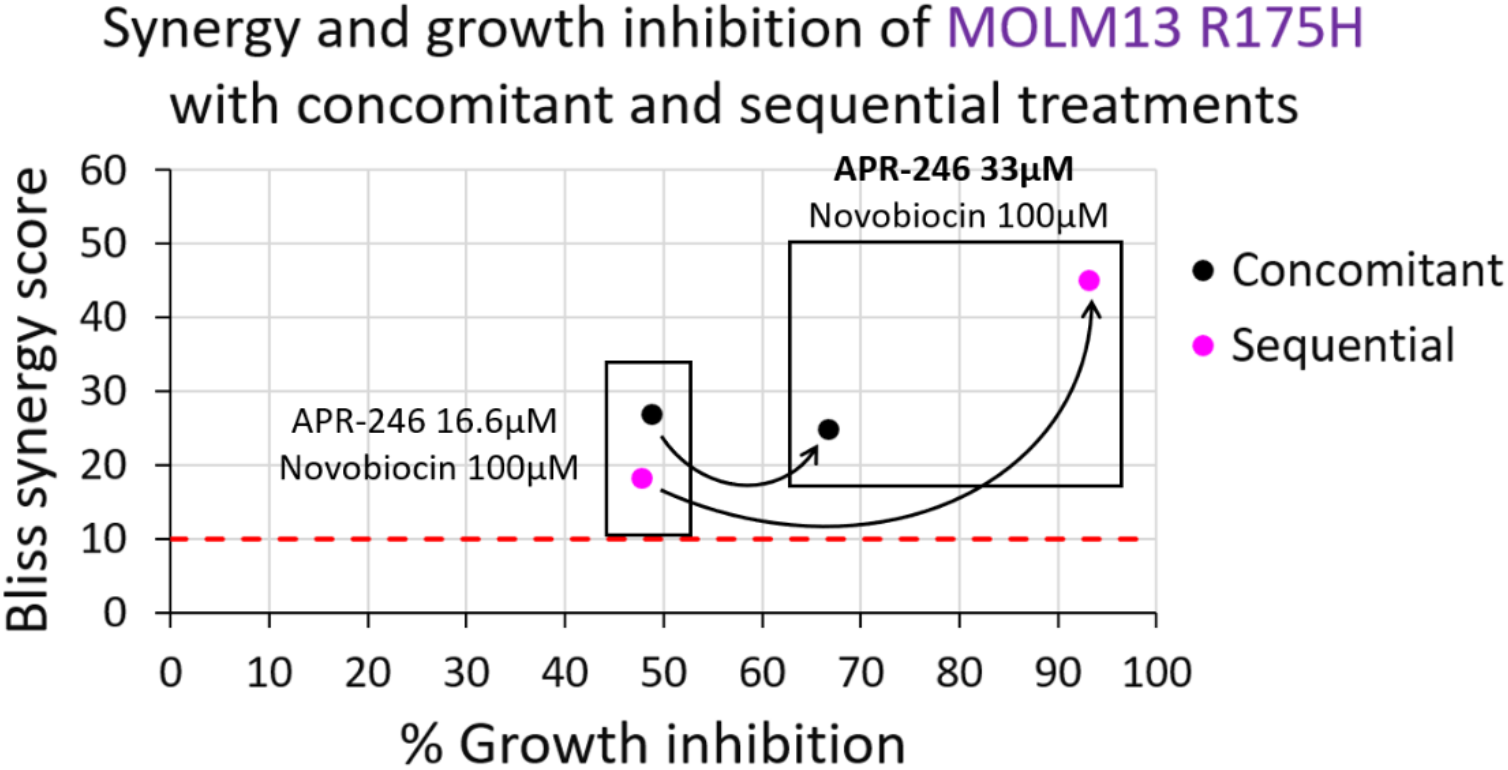
Comparison of concomitant and sequential treatment with APR-246 and NVB in MOLM13 *TP53* R175H cells. (A) Overview of the concomitant and sequential treatment schedules. (B) Comparison of APR-246 monotherapy dose-response curves in different experimental schemes. In the “concomitant treatment” scheme (purple line), APR-246 monotherapy was administered to a cell density of 300,000 cells/mL as described in Fig. 1. In the “concomitant control of the sequential scheme” (black line), cells were seeded at 200,000/mL and APR-246 monotherapy was administered after 24 hours, to a higher cell concentration, of around 350,000 cells/mL. In both experimental schemes viability was assessed 72 hours after incubation with APR-246. APR-246 has higher efficacy when administered to wells with lower cell density (the “concomitant treatment” scheme, purple), with an IC50 of approximately 25 µM (as indicated by the dashed line). (C) Percentage of cell growth inhibition relative to non-treated control following <u>sequential</u> treatment with APR-246 and NVB. (D) Percentage of cell growth inhibition relative to non-treated control following <u>concomitant</u> treatment with APR-246 and NVB. (E) Heatmap representation of Bliss synergy scores for the APR-246 and NVB <u>sequential</u> treatment setup. (F) 3D representation of the Bliss synergy score for the APR-246 and NVB <u>sequential</u> setup. (G) Heatmap representation of Bliss synergy scores for the APR-246 and NVB <u>concomitant</u> treatment setup. (H) 3D representation of the Bliss synergy score for the APR-246 and NVB <u>concomitant</u> setup. The color scale reflects the type of interaction between the two compounds: antagonistic (green), additive (white), or synergistic (red); a synergy score of 10 or above was considered synergistic. The reported data represent an average of two independent replicates. (I) Dot plot comparing the observed percentage of growth inhibition and Bliss synery scores for the sequential (magenta) and concomitant (black) treatment regimens. The differences between combination treatment with APR-246 at 16.6 µM and NVB at 100 µM versus APR-246 at 33 µM and NVB at 100 µM are highlighted.

## Discussion

The compound APR-246 (eprenetapopt) has shown promising results in combination with azacitidine (AZA) in phase II clinical trials in patients with *TP53*-mutant MDS and AML (Cluzeau et al. 2021; Sallman et al. 2021) but a phase III trial in the same indications did not meet study endpoints (aprea.com; Tuval et al. 2024). With the aim of enhancing clinical efficacy of APR-246, we sought to develop novel potent and synergistic combination treatment with APR-246. Here we show that the combination of APR-246 and NVB is synergistic in both a panel of MOLM13 AML cell lines and primary AML/CMML patient samples.

We used the Bliss independence model to calculate synergy scores, based on the fact that the two compounds act on independent molecular pathways. Moreover, unlike other models, Bliss is an effect-based approach and therefore does not require the two drugs to display identical dose-response curves, making it particularly suitable for our experimental setting. We observed the highest synergy in *TP53*-mutant cells with Bliss maximal synergy scores that ranged from 55.12 in cells carrying the R175H *TP53* mutation to 26.87 in those with the R282W mutation. These values are well above the cut-off of 10, confirming potent synergy. The relatively broad range of maximal synergy scores could reflect differences in the expression levels of PolƟ among the cell lines, which influence the extent to which cells depend on this pathway, or the activities of other pathways that contribute to cell detoxification from DNA damage. We also observed synergy in *TP53* WT and null cells, although at a higher concentration of APR-246. This may reflect the reduced sensitivity of these cells to APR-246 as monotherapy. To account for incomplete independent mechanisms of action for the two compounds we calculated ZIP synergy scores which confirmed our findings (Suppl. fig. S4; Suppl. table S5).

Our data suggest that the strong synergism between the two compounds is due to the interaction between the effects of the two compounds. APR-246 generates formaldehyde and NVB inhibits the repair of DNA-protein crosslinks (Fig. 2A) which can increase DNA damage. Despite this, in *TP53*-mutated cells, NVB as monotherapy does not induce apoptosis (Fig. 2B). However, apoptosis can be induced in these cells by the addition of APR-246 (Fig. 2B) which can both target p53 and perturb redox homeostasis (Fig. 3A, B). On the other hand, and to a greater extent, NVB-mediated downregulation of dihydroorotate dehydrogenase (DHODH), a key component of the cellular defence machinery against ferroptosis (Peng et al. 2024), may enhance the known ability of APR-246 to induce oxidative stress and ferroptosis (Birsen et al. 2022; Fujihara et al. 2022). Both of these effects have p53-indpendent components, as confirmed by the observed synergy in *TP53* null cells. However, the enhanced sensitivity to ferroptosis in p53-expressing cells (especially mutant cells) and the presumed reactivation of p53 result in an overall higher effect in *TP53*-mutated cells.

To evaluate the relationship between synergy and efficacy in greater depth, each combination was graphically represented as a function of the corresponding Bliss synergy score and the percentage cell growth inhibition in each MOLM13 subline relative to non-treated controls (Fig. 1 G,H). A Bliss synergy score greater than 10 was considered synergistic and only combinations resulting in at least 50% cell growth inhibition were taken into consideration in terms of efficacy. This graphical analysis enabled the identification of several concentrations of the compounds that demonstrated both significant synergy and high efficacy in suppressing tumor cell growth, highlighting their potential clinical relevance. The combination of APR-246 at 16.6 μM and NVB at 100 μM showed the strongest synergistic effect in all *TP53* mutant cell lines. Importantly, this combination also exhibited good efficacy, exceeding 50% growth inhibition in all mutant cell lines (Fig. 1G). An equivalent level of growth inhibition was observed with APR-246 monotherapy at higher concentrations (at least 25 µM), while NVB as a single agent failed to reach this effect within the investigated concentration range. Thus, the combination treatment proved successful in achieving high efficacy by eliminating at least half of the tumor cells while allowing reduced concentrations of APR-246 and NVB as compared to monotherapies, thus mitigating the associated toxicity. Nevertheless, these plots also emphasize that this combination (APR-246 at 16.6 μM and NVB at 100 μM) does not translate to maximal growth inhibitory effect.

Our cell death analyses showed that combination treatment with APR-246 and NVB induced both ferroptosis and apoptosis the MOLM13 cells. NVB alone increased ferroptosis in all cells, which explains the observed reduced specificity with the combination treatment. APR-246 alone induced a statistically significant increase in ferroptosis compared to NVB alone, and the combination of APR-246 and NVB induced even higher levels of ferroptosis. This increase was more prominent in the *TP53*-mutant and WT cells. The increase in ferroptosis in the mutant cells is mainly driven by the high levels of ferroptosis in R175H and R273H cells, consistent with the differences in synergy scores and efficacy among the cell lines. Thus, our findings indicate that the observed mutant p53-dependent synergy is at least in part mediated by ferroptosis. Altogether, these data suggest that the addition of NVB to APR-246 does not change the mechanism of cell death but rather enhances it, and this effect is more pronounced in *TP53*-mutant cells.

Importantly, we also detected synergy with the combination of APR-246 and NVB in primary AML/CMML patient cells. Consistent with our data from the MOLM13 cell line panel, *TP53*-mutant samples were more sensitive to APR-246 monotherapy with IC50 values in the nanomolar range compared to micromolar range (>15 µM) for the *TP53* wild-type samples. The combination treatment was synergistic in both *TP53* WT and mutant samples but lower concentrations of APR-246 were required for synergy in mutant samples, again in keeping with our results in the MOLM13 cell lines and an overall higher efficacy of APR-246 in *TP53*-mutant cells. The fact that combination of APR-246 at concentrations as low as 0.031 µM with 100 µM NVB achieved potent synergy and 95% grownth inhibition in one *TP53*-mutant sample suggests the possibility that this combination could eliminate *TP53*-mutant leukemic cells in patients with minimal toxicity. The feasibility of the combination treatment for clinical implementation is further underscored by previous clinical data for NVB indicating low toxicity at doses up to 1-2 gr/day which allow a plasma concentration of 100 µM (Eder et al. 1991; Chen et al. 1994).

We also found that sequential administration of APR-246 and NVB could further potentiate their synergistic interaction. Combination of APR-246 at 16 µM and NVB at 100 µM gave higher synergy score in the concomitant treatment (27 vs. 18) and a similar growth inhibition (48%) as compared to the sequential treatment. The sequential treatment achieved its maximal synergy score with the combination of APR-246 at 33.3 µM and NVB at 100 µM; this score was twice as high as the maximal synergy score of the concomitant control, obtained with 16.6 µM APR-246 and 100 µM NVB. Furthermore, when administered sequentially, APR-246 at 33.3 µM and NVB at 100 µm achieved 93% growth inhibition as compared to 66.7% in the concomitant control with the same concentrations. Hence, the sequential administration of NVB showed a highly synergistic interaction and provided significant advantages in terms of cell-killing efficacy compared to the concomitant administration of the same drug concentrations (Fig. 4G).

This study has several limitations. First, assessment of growth inhibition and synergy was performed in the MOLM13 panel of established cell lines, which may not fully reflect the biology of primary MDS/AML patient cells. We have mitigated this by also analyzing several primary patient samples with different *TP53* status. Moreover, we cannot conclude from the DPC analyses that inhibition of repair of DPCs is the critical activity of NVB that accounts for the observed synergies. Also, the fact that MOLM13 cells have high inherent levels of formaldehyde (Vekariya et al. 2023) makes the assessment of DPCs upon combination treatment with APR-246 and NVB more complicated.

In summary, our data demonstrate potent synergy between APR-246 and NVB across the entire panel of MOLM13 AML cells. This synergistic effect is particularly pronounced in p53-expressing cells as compared to *TP53* null cells and most prominent in cells carrying the R175H and R248Q hotspot mutants, indicating that the synergism is at least in part driven by mutant p53-dependent mechanisms. The observed synergy in both *TP53* WT and mutant primary AML/ CMML patient cells suggests that our combination treatment strategy will allow a reduction of the dose of APR-246 while maintaining efficacy and and potentially sparing normal cells carrying WT *TP53*. Further studies will be required to precisely decipher the mechanisms of the observed synergy. Nonetheless, these results suggest that the combination of APR-246 and NVB should be investigated in future clinical trials.

## Materials and methods

### Cells and reagents

MOLM13 is a human leukemia cell line established from the peripheral blood of a 20-year-old male diagnosed with acute myeloid leukemia, FAB subtype M5a. MOLM13 cells were obtained from Prof. Steffen Boetcher, University Hospital of Zürich, Switzerland (Boettcher et al. 2019). Cells had been previously modified using the CRISPR/Cas9 genome-editing technology to introduce the most frequent *TP53* missense mutations, along with null alleles. The specific MOLM-13 cell lines investigated in this work carried bi-allelic alterations in *TP53*. They were hemizygous for the following mutations (carrying one allele with the designated mutation and one allele with a frameshift mutation): R175H, Y220C, M237I, R248Q, R273H, R282W. One cell line carried two wildtype alleles, and one cell line was *TP53*^null^ (both alleles with frameshift mutations). The mutational profile was validated by targeted next-generation sequencing as previously described (Pritzl et al. 2023) (Supplementary table S4). MOLM-13 cells were grown in suspension culture in RPMI-1640 Medium (cat. number R00883-500 mL, Sigma-Aldrich, St. Louis, MO, USA) supplemented with 20% Fetal Bovine Serum (FBS, cat. number A5256801, Gibco, Waltham, MA, USA) heat inactivated at 56°C for 30 minutes and 1% L-Glutamine (cat. number G7513-100 mL, Sigma-Aldrich, St. Louis, MO, USA). Cells were cultured at 0.2-0.4×10^6^ cells/mL in 6-well plates or 10 mL dishes at 37°C, in a humidified atmosphere of 5% CO_2_; they were harvested at their saturation density (< 2.5×10^6^ cells/mL) every three or four days and were counted with Countless II Automated Cell Counter (Thermo Fisher Scientific Waltham, MA USA) after Trypan blue viability staining. APR-246 was provided by Aprea Therapeutics, USA. NVB (Novobiocin) was purchased from Selleck Chem, Houston TX (cat: S2492).

### DNA extraction for sequencing of cell lines

DNA of up to 2×10^6^ cells was extracted using Qiagen Puregene kit (Cat: 158043, Qiagen, Venlo, Netherlands) and was treated with RNAse according to manufacturer’s protocol. DNA quantity and purity were assessed by Nanodrop spectrophotometer (ThermoFisher Scientific Waltham, MA, USA).

### Trypan blue viability assay

For the calibration of the trypan blue cell viability assay, MOLM13 *TP53*^R175H/-^and *TP53*^null^ cells were seeded in 6-well plates and 24-well plates at concentrations ranging from 1×10^5^ to 1×10^6^ cells/mL and were incubated in a humidified atmosphere (+37°C and 5% CO_2_). In order to create growth curves aimed at understanding the optimal culture conditions (seeding and harvest concentrations), cells were counted every day for five days.

### Treatment with APR-246 and NVB

APR-246 was reconstituted in 100% DMSO to obtain a final concentration of 100 mM and was stored in working aliquots at –20°C. NVB sodium at a concentration of 10 mM in DMSO was stored in working aliquots at –80°C. The following concentrations, derived via dilutions, were used for each drug: APR-246 5, 10, 16.6, 25, 33.3, 50 and 75 µM; NVB 25, 50, 75, 100 and 150 µM. For each cell-line, cells in their logarithmic phase of growth were seeded in two 24-well plates at a density of 3×10^5^ cells/mL. Both drugs were directly added to the culture medium on day zero. Each plate included one well with non-treated cells as a control. Cytotoxicity was evaluated after 72 hours by Trypan blue counting. Experiments were repeated at least twice for each cell line. For sequential combination treatment, cell viability was assessed 4 days after seeding. To make sure that the cells are still in their logarithmic growth phase they were seeded on day zero at a concentration of 2×10^5^ cells/mL. NVB at a concentration range of 0-100 µM was added on day zero, whereas APR-246 at concentration range of 0-33.3 µM was added to each well on day 1. As a control (concomitant scheme) both compounds were added to the cell culture on day 1. Cell viability was assessed 72 hours after treatment with APR-246 (sequential scheme) or with the combination APR-246 and NVB (concomitant scheme). Standard and sequential treatment with APR-246 and NVB were repeated twice for each cell line. Data was analysed using *SynergyFinder Plus* R package (Zheng et al., 2022) with default parameters unless specified otherwise. Results from the two replicates were combined and analyzed together.

Missing values in the dose-response matrix (specifically, for the combination of APR-246 75 µM and NVB 150 µM) were automatically imputed by the *mice* R package. To estimate the synergistic effects of a drug combination, reference models assume that the drug’s effect is expressed as a continuous value between 0% and 100%; nevertheless, negative readouts can occur in practice, potentially introducing technical errors in the dose-response data. To avoid this, as suggested by the package-user tutorial, a baseline correction for negative inhibition values was applied on the raw dose-response data. The *SynergyFinder Plus* package calculates synergy scores according to four of the major reference models, including HSA, Loewe, ZIP and Bliss. To calculate the synergy scores for the experiments presented in this work we employed the Bliss independence model (Bliss 1939), as its underlying assumptions best suited the characteristics of the two drugs. Unlike models that require fitting parametric curves for the two single treatments (monotherapies), the Bliss model is an effect-based approach and utilizes only the raw monotherapy dose-response data to calculate synergy, under the assumption of non-interaction (Vlot et al., 2019). Synergy was confirmed by calculating the ZIP synergy score that detects synergy if the dose-response curve of the combination treatment is different than that of each individual drug (Yadav et al., 2015). We did not use the Loewe score because its basic assumption is that for a given effect of drug A, an equivalent dose for drug B exists. The low inhibitory effect of NVB as monotherapy, where IC50 was not achieved, violates this assumption. Dose response curves for monotherapies were generated by a 3-parameter log-logistic model using the *drc* R package (Ritz et al, 2015).

### DPC assay

Cells (with at least 90% viability) were seeded at 300,000 cells/mL in 6 well plates. Cells were supplemented with media (“not-treated”) or one of the following treatments: NVB, APR-246, formaldehyde (ThermoFisher Scientific Waltham MA USA, Cat: 28906) or their combination at the designated concentrations, gently mixed and incubated for 6 hours. Cells were then harvested and processed according to the advanced recovery of K (potassium)-SDS precipitates (ARK) protocol (Hu et al. 2020). DNA fractions were measured with Qubit fluorometer (High sensitivity kit, ThermoFisher Scientific Waltham MA USA, Cat: Q33230). Experiments were performed in duplicates (cells from a single culture were split into two wells and incubated and processed independently) and repeated twice (two biological replicates). The mean of the two technical replicates within each experiment was used as the value that represents that experiment.

### Cell death assays

Cells (with at least 90% viability) were seeded at 300,000 cells/mL in 24 well plates. Cells were supplemented with media (“not-treated”) or with APR-246, NVB or with their combination in the designated concentrations. Each biological replicate was generated by partitioning a single parent cell culture on a given day, and at least three biological replicates were performed for each cell line. BODIPY C11 (Image-iT™ Lipid Peroxidation Kit, ThermoFisher Scientific Waltham MA USA, Cat: C10445) was diluted 1:10 in Dulbeccoʹs Phosphate Buffered Saline (PBS, Cat: 59321C, Sigma Aldrich, Saint Louis, MO, USA) to a final concentration of 1 mM. It was then added to each well (apart from well that were used as unstained control) to achieve a final concentration of 2 μM. Wells were gently mixed and incubated for 30 minutes at 37°C in the dark. Cells were harvested, washed twice with 2mL PBS and counted. Cells were then resuspended in Annexin V binding buffer 10X (Cat: 556454, BD Biosciences, San Jose, California, USA) that was pre-diluted 1:10 with distilled water and brought to room temperature to achieve 1×10^6^ cells/mL. 100 μL (100,000 cells) were transferred into a new tube where they were mixed with 5 μL of fluorochrome-conjugated (V450) Annexin V (Cat: 560506, BD biosciences, San Jose, California, USA) and Propidium Iodide (PI, Cat: P4864, Sigma Aldrich, Saint Louis, MO, USA) in a final concentration of 50 ngr/100,000 cells in 100 μL. Cells were incubated for 15 minutes in room temperature in the dark. 400 μL of “Annexin V binding buffer x1” were added to each tube samples were recorded by NovoCyte Quanteon 4025 flow cytometer (Agilent Technologies, Santa Clara, California USA). Data were analyzed using NovoExpress software (version 2.2.0). Gating strategy is shown in Supplementary Figure S2A. PE and FITC intensities (for assessment of ferroptosis) were recorded only for PI-negative viable cells to avoid analyzing cells that have lost membrane integrity, which could lead to non-specific oxidation of BODIPY C11 and that can artificially elevate the FITC signal. Therefore, cells with side scatter and forward scatter (FSC) properties that are characteristic of small or dying cells (“Cells 2”, Fig. S2A) were excluded from analysis of ferroptosis (but were included in the analysis of apoptosis).

Identical PI threshold values were used for both the apoptosis and the ferroptosis analyses. At least 20-30,000 cells were included in the analysis of each sample. For analysis of ferroptosis, the ratio between the measured fluorescence at 590 nm (PE channel) and the measured fluorescence at 510 nm (FITC) were calculated for each individual cells and presented as density plot. To quantify the percentage of ferroptotic cells in each biological replicate, we used as a cut-off the PE to FITC ratio value of the 5^th^ percentile in the non-treated sample of that replicate (dashed blue line in Fig. 2C and D). Cells that were pre-incubated for 2 hours at 37°C with 100uM of Cumene hydroperoxide (included in the Image-iT™ Lipid Peroxidation Kit) were used as positive control for calibration to determine the final staining concentration of the BODIPY C11 reagent (2μM) as well as to generate the compensation matrix that was used in all the experiments to account for fluorescence spectral overlap and spillover.

### Patient samples

Primary bone marrow and/or peripheral blood were obtained from the Patnaik lab patient consented Biobank under Mayo regulatory guidelines (the Mayo Clinic Clonal Hematopoiesis/ MPN biobanks IRB 16-004173 and 11-005599). Mononuclear cells of patients with chronic myelomonocytic leukemia or AML with or without a pathogenic TP53 mutation were enriched using density gradient centrifugation. Mutational status was determined by clinical NGS panel (Pritzl et al., 2023).

### Colony forming assays (CFA)

Briefly, approximately 80,000 cells were plated in human recombinant cytokine supported methylcellulose (H4434 MethoCult, StemCell Technologies, Canada). Concentrations of APR-246 and Novobiocin were added and cells were incubated for 10 days at 37°C with 5% CO2. At day 10, colonies were qualitatively assessed using light microscopy for size and type (BFU-E = erythroid, GM = granulocyte/monocyte mixed, G = granulocyte, GEMM = primitive mixed erythroid/myeloid). IC50 was generated by Graph Pad Prism software version 10, using normalized response non-linear regression.

### CellGlo proliferation assay

Cells were plated in 96 well format using combinations of APR-246 and Novobiocin. Each drug concentration or combination was plated in at least triplicates. For the assay, 12,000-20,000 cells were plated according to the commercial kit protocol: CellTiter-Glo Luminescent Cell Viability Assay (Promega), and read out via luminometer. Percent viability was calculated and SynergyFinder 3.0 (Ianevski et al., 2022) software was used to calculate percentage of inhibition and Bliss synergy scores. For monotherapy IC50 values were calculated via dose-response four parameter nonlinear regression by Prism software.

### Data analysis

Statistical analyses (as described for each section) and plots were performed in R (version 4.6.1), using appropriate packages.

### Statistical analyses of ferroptosis and apoptosis flow cytometry data

Data were analyzed using a non-parametric split-sample, repeated-measures, two-way-mixed model ANOVA-type design. Since the treatment wells were generated by partitioning a single parent cell culture on a given day, observations within the same experimental day are statistically non-independent. To account for this dependent structure (repeated measures structure), the day of the independent experiment was treated as a random blocking factor. This approach controls for the shared baseline environmental variance inherent to cell cultures harvested and treated concurrently on the same day. The percentage of ferroptotic or apoptotic cells that were measured in each cell line’s non-treated wells in each experimental day were subtracted from the percentages that were measured on that experimental day in treated wells of that cell line. Prior to formal hypothesis testing, statistical assumptions were evaluated by extracting the residuals from a linear model that included the main effects of cell type and treatment, while controlling for the experimental day as a blocking factor. Due to differences in baseline percentage values across cell lines, raw residuals were converted to standardized residuals to eliminate scale-dependent artifacts. The variances of the standardized residuals across the experimental groups ranged from 0.17 to 5.37 (a 31-fold difference), violating the homoscedasticity assumption required for traditional parametric ANOVA. Thus, an ANOVA-Type Statistic (ATS) for an F1-LD-F1 design (with a unique subject identifier combining cell line and experimental day) was employed to accommodate both the unequal group variances and the repeated-measures split-sample design (Brunner et al 1997) using the nparLD package in R (Noguchi et al 2012). Hypothesis testing was explicitly evaluated under the unweighted relative marginal effects framework, which acts as a non-parametric Behrens-Fisher adjustment.

For analysis of apoptosis data and determining which cell line-treatment combinations drive the interaction, planned, a priori, interaction contrasts (including difference-of-differences tests) were constructed according to data visualization and evaluated simultaneously using non-parametric multiple contrast test via the nparcomp package in R (Konietschke et al. 2015). To accommodate the small sample size and group heteroscedasticity within the split-sample design, a multivariate t-distribution with a Satterthwaite approximation was utilized to calculate data-driven, fractional degrees of freedom. Multiple comparisons were controlled simultaneously using a joint multivariate single-step max-t procedure to maintain the family-wise error rate at α = 0.05.

For analysis of ferroptosis data, post hoc analysis for pairwise comparisons for the treatment main effect were performed with Kenward-Roger residual degrees of freedom (to adjust for the small-sample repeated measures design) and p-values were adjusted using Bonferroni’s correction for multiple comparisons. For the main effect of cell line over ferroptosis, post hoc analysis of seven planned custom contrasts was performed. For each of these seven planned contrasts, the raw biological effect size was calculated as a weighted linear combination of the cell line marginal means [ΣwiX ̅i] where wi represents the directional weights for each custom contrast, and X ̅i represents the unweighted average of the individual cell-treatment combination group means for cell line i. The descriptive standard errors of the mean (SEMs) for these absolute percentage differences were calculated using standard linear combination variance propagation [Var(ΣwiX ̅i) = Σwi2Var(X ̅i)/nt], where Var(X ̅i) is the variance of X ̅i and nt is the number of collapsed treatment conditions (nt = 3) (Hogg et al., 2019). These calculations were required since standard errors cannot be directly averaged or divided when combining asymmetrical, weighted groups. For the relative fold-change ratios, the numerator (target pooled mean) and denominator (comparison pooled mean) were isolated according to the directional weights of the contrast vectors. The corresponding descriptive standard errors for these ratios were propagated using first-order Taylor series expansion via the Delta method (Geary’s approximation), which accounts for the interacting uncertainties of both the shifting numerator and denominator variables (Casella et al. 2002).

Standardized effect sizes for the seven planned custom cell line contrasts were calculated as standardized contrast metrics (Cohen’s d equivalents) via the emmeans package in R. Effects were standardized by dividing the weighted contrast estimate by the linear mixed-effects model’s residual standard error to account for the pooled residual variance across the multigroup framework. To optimize statistical power while maintaining rigorous error control across the seven custom hypotheses, p-values were adjusted simultaneously for all seven contrasts using the Benjamini-Hochberg False Discovery Rate (FDR) procedure (Benjamini et al. 1995). Descriptive values for treatment and cell-type effects are reported in text and figures as raw percentage data (Mean ± SEM) rather than arbitrary model ranks. Standardized effect sizes (Cohen’s d) and significance testing were strictly computed on the aligned ranks to satisfy non-parametric assumptions and were adjusted to account for the random block effect of experimental day. All statistical tests were two-tailed, and effects were considered statistically significant at an adjusted p-value of < 0.05. All analyses were executed in R (version 4.6.1).

## Acknowledgements

This work was supported by grants to KGW from Radiumhemmets Forskningsfonder (251461), the Swedish Cancer Society (24 3630 Pj 01 H), the Swedish Research Council (VR 2021-02064), and Karolinska Institutet, and by grants to MMP from Kura Oncology, Stem Line Pharma, Epigenetix, Polaris and Solu Therapeutcis. We thank Prof. Steffen Boettcher, University of Zürich, for kindly providing the MOLM13 cell line panel.

