## Supplementary material for "APR-246 and DNA polymerase theta inhibitor Novobiocin synergize to eliminate *TP53* mutant AML cells": Suppl figs and tables

**Tuval et al. Supplementary Figures and Tables**

**Suppl. Fig. S1A**

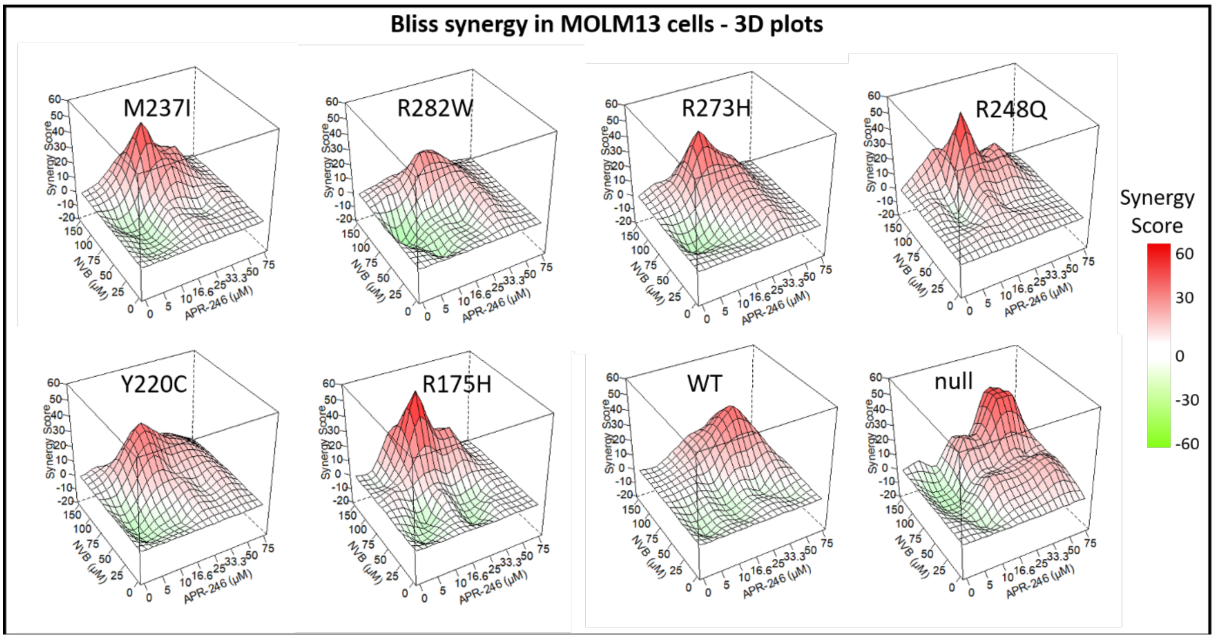


3D plots for the Bliss synergy scores obtained for each of the cell lines. Data were generated by two biological replicates.

**Suppl. Fig. S1B**

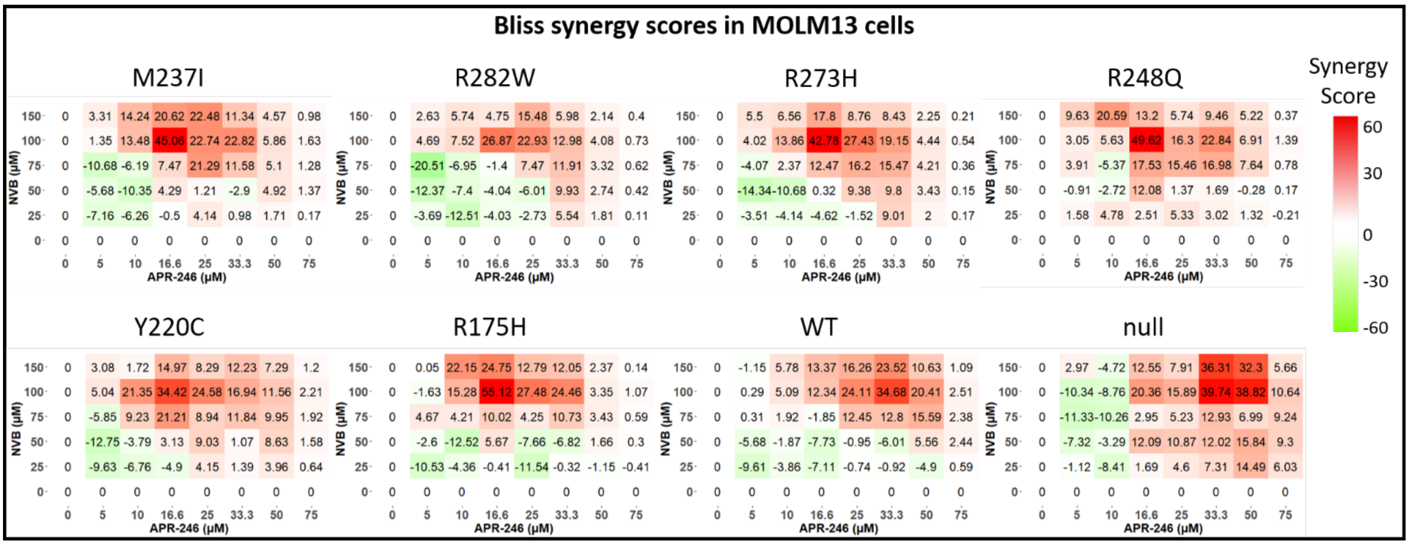


Same data as in Fig. S1A, presented as heatmap plots with the Bliss synergy scores for each drug combination.

**Suppl. Fig. S1C**

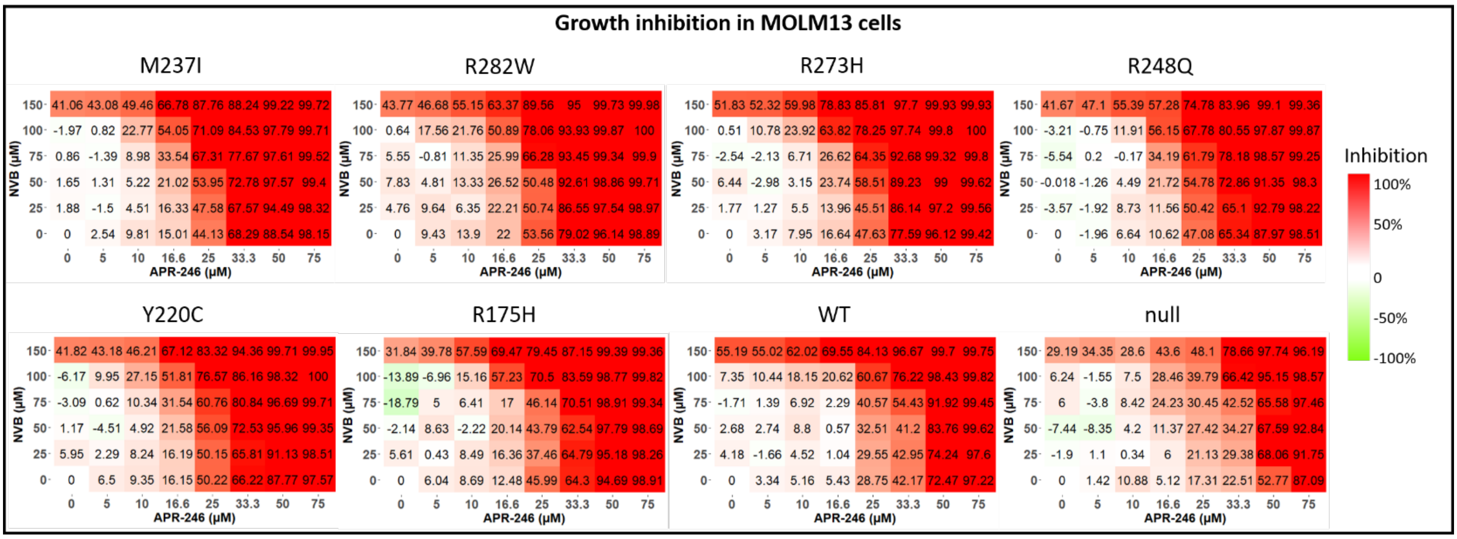


Heatmap plots for the percentage of growth inhibition induced by each drug combination for each of the cell lines. Average values of the normalized (to non-treated cells) percentage of growth inhibition in two biological replicates.

**Suppl. Fig. S1D**

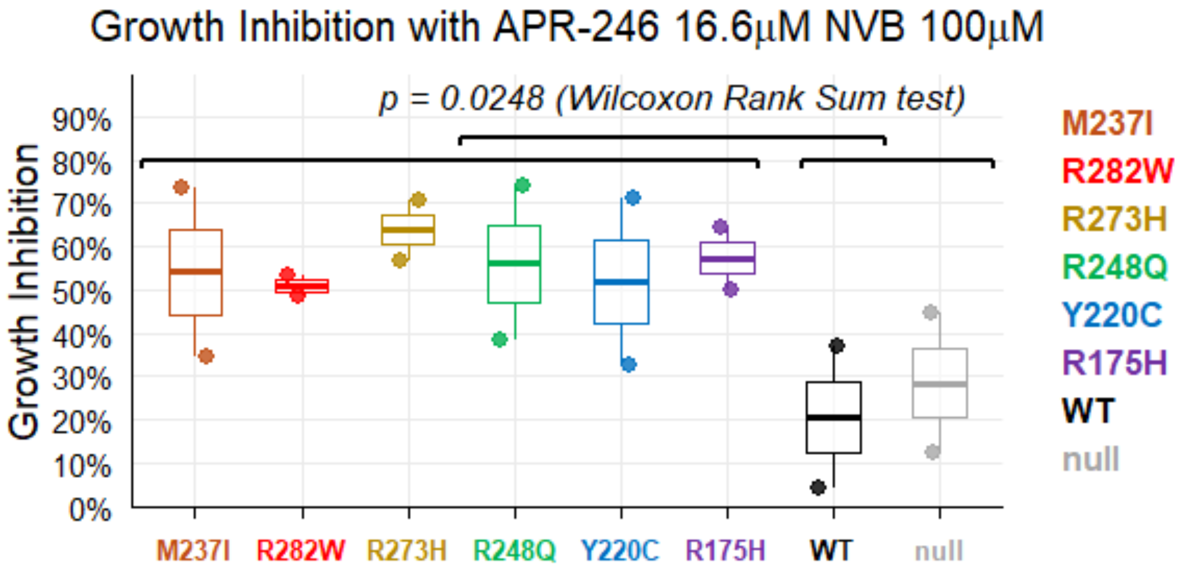


Box plot for the percentage of growth inhibition across all tested MOLM13 cells. Each dot represents a biological replicate. Boxes represent the interquartile range (IQR) and medians are shown as bars. Whiskers represent values less than the 25th percentile and greater than the 75th percentiles, and they extend no more than 1.5 × IQR units away from either end of the boxes.

**Suppl. Fig. S2**

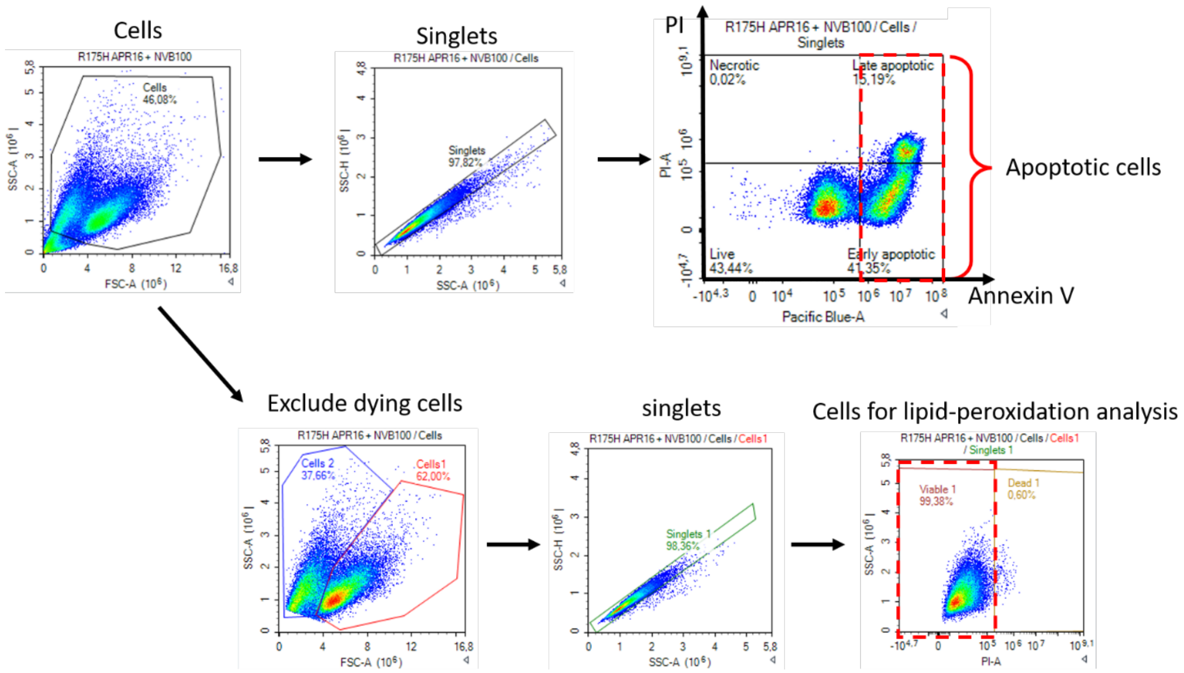


Representative flow cytometry plots.

<u>Upper panel: Gating strategy for measurement of apoptotic cells.</u>

Singlets (identified by a linear correlation between their side scatter height (SSC-H) and side scatter area (SSC-A) values, were divided according to PI positivity and Annexin V positivity:
PI-negative, Annexin V-negative cells were regarded as “Live”
PI-negative, Annexin V-positive cells were regarded as “Early apoptotic”
PI-positive, Annexin V-positive cells were regarded as “Late apoptotic”
PI-positive, Annexin V-negative cells were regarded as “Necrotic”.
Early and late apoptotic cells were combined and considered as “apoptotic”.

<u>Lower panel: Gating strategy for assessment of lipid-peroxidation.</u>

PE and FITC intensities (for assessment of ferroptosis) were recorded only for PI-negative viable cells to avoid analyzing cells that have lost membrane integrity, which could lead to non-specific oxidation of BODIPY C11 and that can artificially elevate the FITC signal.
Cells with side scatter and forward scatter (FSC) properties that are characteristic of small or dying cells (“Cells 2”, blue) were excluded from analysis of ferroptosis (but were included in the analysis of apoptosis). Singlets (identified by a linear correlation between their SSC-H and SSC-A) out of “Cells 1”, were divided according to PI positivity. PI-negative cells (“Viable 1”) were included in the analyses.

Identical PI threshold values were used for both the apoptosis and the ferroptosis analyses.

**Suppl. Fig. S3A**

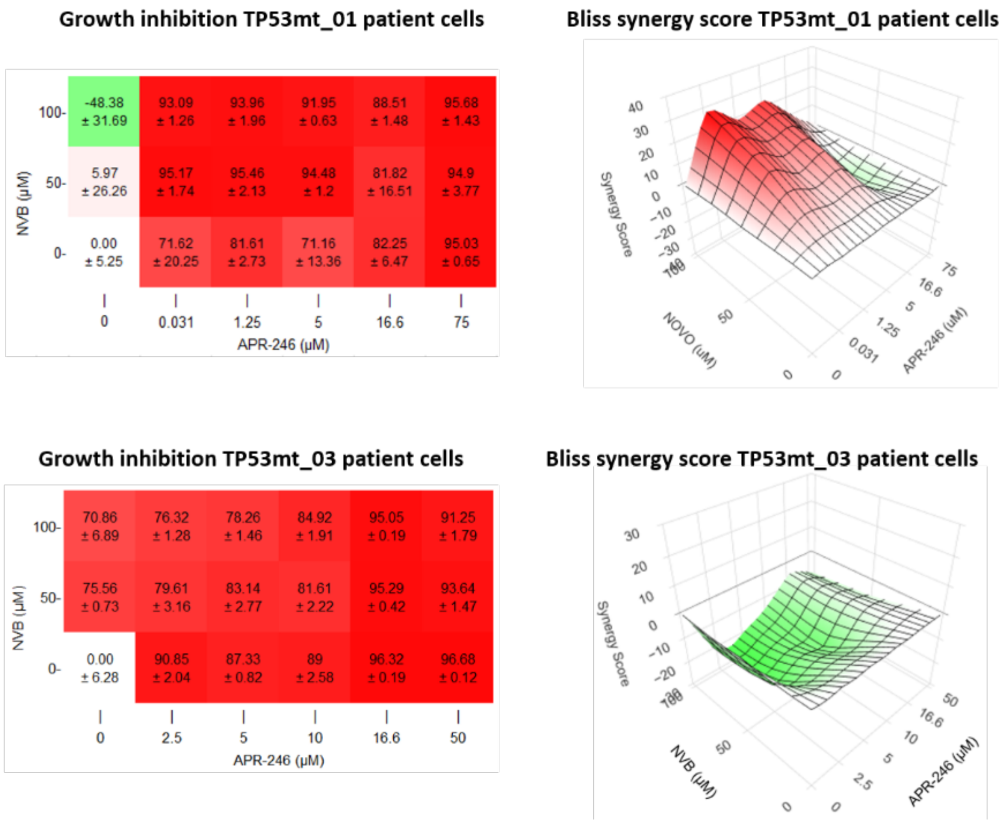


Growth inhibition and Bliss synergy scores in *TP53*-mutated primary peripheral blood or bone marrow samples from Mayo Clinic patients with CMML and/or AML.

Left panels: Heatmap plots for the growth inhibition induced by each drug combination for each of the samples. The indicated values are the normalized (to non-treated cells) values of the average of at least 3 replicates.

Right panels: 3D plots for the Bliss synergy scores obtained for each of the samples.

**Suppl. Fig. S3B**

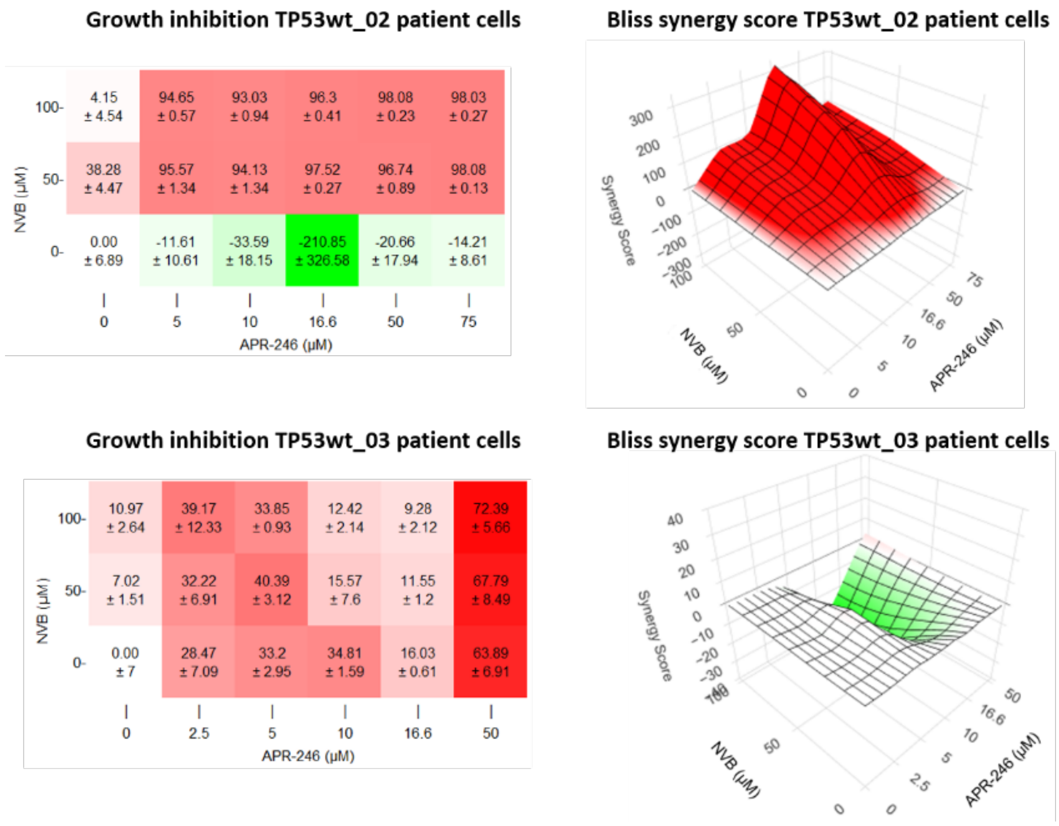


Growth inhibition and Bliss synergy scores in *TP53*-wild type primary peripheral blood or bone marrow samples from Mayo Clinic patients with CMML.
Left panels: Heatmap plots for the growth inhibition induced by each drug combination for each of the samples. The indicated values are the normalized (to non-treated cells) values of the average of at least 3 replicates.
Right panels: 3D plots for the Bliss synergy scores obtained for each of the samples.

**Suppl. Fig. S4**

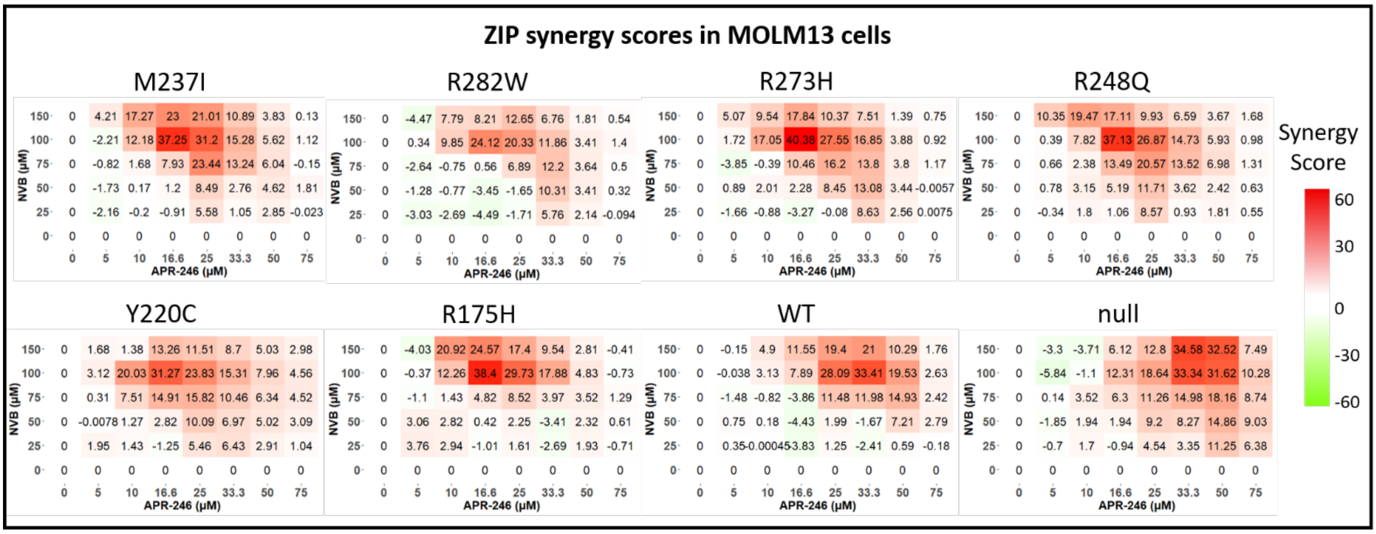


Heatmap plots for the ZIP synergy scores obtained for each of the cell lines. Data were generated by two biological replicates.

**Suppl. Table S1**

Growth inhibition and Bliss synergy scores obtained with APR 16.6µM and NVB 100µM

| Cell line | % Growth inhibition | Synergy score |
|---|---|---|
| R175H | 57.23 | 55.12 |
| R273H | 63.82 | 42.78 |
| R282W | 50.89 | 26.87 |
| Y220C | 51.81 | 34.42 |
| M237I | 54.05 | 45.06 |
| R248Q | 56.15 | 49.62 |
| WT | 20.62 | 12.34 |
| null | 28.46 | 20.36 |

**Suppl. Table S2**

Growth inhibition and Bliss synergy scores obtained with APR 33.3µM and NVB 100µM

| Cell line | % Growth inhibition | Synergy score |
|---|---|---|
| R175H | 83.59 | 24.46 |
| R273H | 97.74 | 19.15 |
| R282W | 93.93 | 12.98 |
| Y220C | 86.16 | 16.94 |
| M237I | 84.53 | 22.82 |
| R248Q | 80.55 | 22.84 |
| WT | 76.22 | 34.68 |
| null | 66.42 | 39.74 |

**Suppl. Table S3A**

Effect size, fold change and statistics of *post hoc* comparisons for treatment main effect over ferroptosis

| comparison | adj. p-value (Bonferroni) | Fold Change | Cohen's *d* |
|---|---|---|---|
| APR-246 16.6µM vs. NVB 100µM | <0.0001 | 5.17 | 1.71 |
| APR-246 16.6µM and NVB 100µM vs. NVB 100µM | <0.0001 | 12.67 | 2.88 |
| APR-246 16.6µM and NVB 100µM vs. APR-246 16.6µM | 0.0058 | 2.45 | 1.17 |

**Suppl. Table S3B**

Effect size, fold change and statistics of seven planned *post hoc* comparisons for cell line main effect over ferroptosis

| comparison | adj. p-value (BH) | Fold Change | Cohen's *d* |
|---|---|---|---|
| p53-expressing vs. null | 0.044 | 3.00 | 1.03 |
| mutants vs. null | 0.044 | 3.18 | 1.05 |
| mutants vs. WT | 0.866 | 1.29 | 0.08 |
| R175H vs. null | 0.044 | 4.20 | 1.50 |
| R273H vs. null | 0.044 | 3.24 | 1.12 |
| WT vs. null | 0.044 | 2.47 | 0.97 |
| R282W vs. null | 0.425 | 2.11 | 0.52 |

p-values were adjusted simultaneously for all seven contrasts using the Benjamini-Hochberg False Discovery Rate (FDR) procedure (Benjamini et al., 1995).

**Suppl. Table S4**

| Sample Name | Gene name | Nucleotide Change | Protein Change | VAF | Consequence |
|---|---|---|---|---|---|
| MOLM13 WT | *TP53* | no variant found | | | |
| MOLM13 R273H | *TP53* | c.215C>G | P72R | 48.26% | missense_variant |
| | | c.794_795del | L265Rfs*6 | 49.43% | frameshift_variant |
| | | c.818G>A | R273H | 47.44% | missense_variant |
| MOLM13 Y220C | *TP53* | c.215C>G | P72R | 49.01% | missense_variant |
| | | c.643A>T | S215C | 47.87% | missense_variant |
| | | c.659A>G | Y220C | 43.34% | missense_variant |
| | | c.659del | Y220Lfs*27 | 48.10% | frameshift_variant |
| MOLM13 M237I | *TP53* | c.215C>G | P72R | 46.15% | missense_variant |
| | | c.711G>A | M237I | 48.30% | missense_variant |
| | | c.728_729insTT | M243Ifs*5 | 50.57% | frameshift_variant |
| MOLM13 R248Q | *TP53* | c.215C>G | P72R | 41.47% | missense_variant |
| | | c.742_743insAGAC | R248Qfs*17 | 48.89% | frameshift_variant |
| | | c.743G>A | R248Q | 45.56% | missense_variant |
| MOLM13 R175H | *TP53* | c.215C>G | P72R | 43.41% | missense_variant |
| | | c.508_509insATGC | T170Nfs*12 | 55.50% | frameshift_variant |
| | | c.524G>A | R175H | 46.15% | missense_variant |
| MOLM13 R282W | *TP53* | c.844C>T | R282W | 42.90% | missense_variant |
| | | c.831_835del | C277Wfs*27 | 56.23% | frameshift_variant |
| | | c.215C>G | P72R | 46.76% | missense_variant |
| MOLM13 null | *TP53* | c.567_577del | P190Sfs*15 | 35.29% | frameshift_variant |
| | | c.572dup | Q192Sfs*17 | 99.08% | frameshift_variant |
| | | c.215C>G | P72R | 38.08% | missense_variant |

Results of targeted next generation sequencing for validation of the mutational profile of the MOLM13 cell lines used in this work. Sequencing was performed as described by Pritzl et al. 2023. VAF, variant allele frequency.

**Suppl. Table S5**

| Cell line | Maximal ZIP score | Combinatorial drug concentrations that achieved maximal ZIP synergy score | |
|---|---|---|---|
| | | APR-246 (μM) | NVB (μM) |
| M237I | 37.25 | 16.6 | 100 |
| R282W | 24.12 | 16.6 | 100 |
| R273H | 40.38 | 16.6 | 100 |
| R248Q | 37.13 | 16.6 | 100 |
| Y220C | 31.27 | 16.6 | 100 |
| R175H | 38.4 | 16.6 | 100 |
| WT | 33.41 | 33.3 | 100 |
| null | 34.58 | 33.3 | 150 |

Maximal ZIP synergy scores and the drug combinations that achieved them in MOLM13 cells.
